# *In silico* reconstruction of a salmonid alphavirus virion reveals distinctive structural and molecular features implicated in virulence *in vivo*

**DOI:** 10.1101/2025.09.19.677251

**Authors:** Stéphane Biacchesi, Calvin Fauvet, Emilie Mérour, Julie Bernard, Annie Lamoureux, Delphine Lallias, Jean K. Millet

## Abstract

Salmonid alphavirus (SAV) poses a significant disease threat to aquaculture. Recently, new alphaviruses from several fish species have been discovered. However, little is known about their biology and diseases potential. Alphaviruses are considered to have originated from a marine environment; therefore, studying fish alphaviruses can inform on the evolutionary history of the genus. Contrary to many terrestrial alphaviruses, there are currently no experimentally-determined structures for aquatic alphaviruses, severely limiting their study. In this work, we harness the power of structural bioinformatics and AlphaFold to reconstruct an entire SAV virion, thereby revealing an exposed and distinctive α-helical feature in its E2 envelope protein. Using an integrative approach, we explore the sequence diversity and evolutionary conservation of this predicted feature and investigate the functional consequences of variations on viral fitness and virulence. This study provides a novel framework paving the way to better understand aquatic alphavirus pathogenicity and host species adaptation.

## Introduction

Epizootics caused by emerging and re-emerging viruses are a serious threat to the development of aquaculture which is recognized as the fastest-growing food sector. Salmonid alphavirus (SAV) is a WOAH-listed viral pathogen that causes sleeping disease (SD) in rainbow trout (*Oncorhynchus mykiss*) and pancreas disease in Atlantic salmon (*Salmo salar*) ^1^.

Members of the *Alphavirus* genus within the *Togaviridae* family are RNA viruses found in diverse aquatic and terrestrial ecosystems and infect invertebrates, fishes, birds, and mammals. Most terrestrial alphaviruses are arboviruses posing a significant threat for human and animal health. A landmark genome-wide study suggested that alphaviruses originated from a marine environment ^2^. This was confirmed by a large scale metatranscriptomics study, which demonstrated that RNA viruses infecting fish species, such as alphaviruses and rhabdoviruses, form lineages basal to those associated with vector-borne terrestrial viruses ^3^. Recently discovered alphaviruses include viral species infecting several fish hosts such as striated frogfish (*Antennarius striatus*), crested flounder (*Plagiopsetta sp.*), jawless inshore hagfish (*Eptatretus burgeri*), and Mediterranean comber (*Serranus cabrilla*) ^3,4^. Whether these viruses found in wild reservoir species can pose a disease threat to aquaculture species remains to be addressed. Despite these advances in virus discovery, little is known about their biology, the disease they cause, and their relationship to SAV and terrestrial alphaviruses.

SAV is the best-studied aquatic alphavirus and like other members of its genus it has a positive-sense non-segmented RNA genome (11-12 kb range) that is 5’ capped and 3’ polyadenylated. Its genome encodes four non-structural proteins (nsP1-4) which are first synthesized as a polyprotein which is then proteolytically processed. Alphavirus nsP proteins mediate viral transcription, replication, and counteract host innate immune responses. A subgenomic RNA encodes the structural polyprotein composed of capsid (Cp), E3, E2, 6k, and E1 proteins, which are also cleaved into individual proteins by proteolytic enzymes. As their name implies, the structural proteins participate in virion formation. There are six major subtypes of SAV, numbered 1 to 6, based on partial sequences of nsP3 and E2. Distinctive SAV features setting them apart from terrestrial alphaviruses include shorter non-coding regions at the 5’ and 3’ genomic ends, a low temperature of replication (10-15 °C), and transmission occurring directly through water without the need for a vector, although SAV nucleic acids have been detected in sea lice (*Lepeophtheirus salmonis*) ^5–7^. Another unique characteristic of SAV is the longer length of the E1 and E2 envelope proteins, due to the presence of relatively long (∼10 aa) inserts. The role of these inserts and more broadly the structure-function relationships of fish alphavirus proteins constitute major limitations in our understanding of their biology and evolutionary relationship with their terrestrial counterparts.

Much of what is known about alphavirus structure stems from studies on terrestrial representatives such as Chikungunya virus (CHIKV) ^8^ and Venezuelan equine encephalitis virus (VEEV) ^9^. Although enveloped, the surface of alphavirus virions (∼70 nm) adopts an ordered *T* = 4 icosahedral symmetry architecture composed of 80 trimeric spikes with each trimer made of heterodimers of E2 and E1, which are both transmembrane proteins and the main determinants of host cell receptor attachment and membrane fusion, respectively ^10^. Within the alphavirus virion, the Cp protein encapsidates the RNA genome forming an inner nucleocapsid shell ^9^. To date, there are no available experimentally-determined structures for SAV or any other fish alphavirus representatives. The lack of widely accessible structural data limits the depth of structure-function analyses of these viral proteins. Since E1 and E2 proteins are the main drivers of viral entry and host range, the absence of structural information precludes a deeper understanding of the molecular determinants of virulence, host specificity, and interspecies transmission.

This is of particular importance for SAV because single amino acid substitutions within the E2 envelope protein were shown by our group to profoundly impact virulence in rainbow trout, mirroring similar phenomena observed with terrestrial alphaviruses ^11^. A similar attenuation was described for CHIKV strain 181/clone 25, a live-attenuated vaccine candidate derived from a South-East Asian human isolate, where only two E2 mutations in domains responsible for receptor recognition were found to alter virulence^12^. A single substitution (A226V) in CHIKV E1 fusion protein identified in an epidemic strain from La Réunion island was shown to be responsible for a significant increase in infectivity in *Aedes albopictus*, providing a plausible explanation for how this variant caused an epidemic in a region lacking its typical vector, *A. aegypti* ^13^. Moreover, our group has previously demonstrated that single substitutions within an epitope motif of SAV2 E2 were associated with altered virulence *in vivo* ^14^. Similar mutational sensitivity (*i.e.* small amino acid changes in envelope proteins leading to large and measurable phenotypical effects) has also been described for other alphaviruses, including VEEV, Sindbis virus (SINV), and Semliki Forest Virus (SFV) ^15–17^. The findings that such mutational sensitivity is shared among several terrestrial and aquatic alphaviruses suggest it could be a genus-wide conserved trait. Since SAV occupies a basal position within *Alphavirus* genus phylogeny, its study can not only inform the biology of fish alphaviruses but also the evolutionary relationships and commonalities with terrestrial alphaviruses.

Recently, structural bioinformatics has undergone a revolution with the emergence of artificial intelligence-based algorithms capable of highly accurate protein structure predictions, as exemplified by AlphaFold ^18,19^. The development of these tools is particularly useful for virology as, until recently, a huge proportion of viral protein sequences had no corresponding structural information. Importantly, these predictive programs open up the possibility to conduct structure-guided studies on viral proteins that previously lacked experimentally-obtained structures.

Here, we leverage the groundbreaking advances brought by AlphaFold and other structural bioinformatics tools by modeling the structural proteins of SAV and reconstructing the structure of a complete virion. Using an integrative strategy, we uncover a unique α-helical structural feature found at a site near the N-terminus (N-term) of E2 that is shared among aquatic alphaviruses. By combining phylogenetic analysis, reverse genetics and *in vitro* and *in vivo* assays, we delve into the sequence diversity of a broad set of SAV isolates to investigate how variations found at this site can impact replicative fitness and virulence in rainbow trout. This work will allow to better anticipate the functional impacts of virulence-associated mutations, which may inform surveillance strategies for disease outbreak investigation and ultimately offer novel insights into the evolutionary dynamics of alphaviruses.

## Results

### Phylogeny of SAV structural proteins

To position SAV and more recently identified fish alphaviruses within the context of the *Alphavirus* genus, the sequences of structural proteins of 17 alphaviruses were aligned to generate a Maximum-Likelihood (ML) phylogenetic tree of representatives from both aquatic and terrestrial environments (Figure 1A & Table S1). The overall topology recapitulates previously established phylogenetic relationships among the *Alphavirus* genus and highlights the branching of fish-infecting alphaviruses forming a distinct basal clade from that of the other alphaviruses. This analysis is in line with the evidence for the ancient evolutionary history of Alphaviruses and their proposed marine origin ^2,3^. Notably, aquatic alphaviruses that infect mammalian hosts, such as Southern elephant seal virus (SESV) and Alaskan harbor porpoise alphavirus (AHPV) are more closely related to terrestrial alphaviruses from the Semliki Forest Virus (SFV) antigenic complex group, with a solidly-supported branching (bootstrap support of 97). Among fish-infecting aquatic alphaviruses, most branch nodes are strongly supported (bootstrap support of 100), with only one node with a relatively lower value (bootstrap support of 63) corresponding to the branching between salmonid alphavirus (SAV, ICTV species *Alphavirus salmon*) and Wenling hagfish alphavirus (WHAV), which is most closely related to SAV. SAV infects Atlantic salmon (*Salmo salar*), rainbow trout (*Oncorhynchus mykiss*), Arctic charr (*Salvelinus alpinus*) and common dab (*Limanda limanda*). Among species closely related to SAV, WHAV was isolated from inshore hagfish (*Eptatretus burgeri*), and is followed by Comber alphavirus (CAV) which was isolated from *Serranus cabrilla*. A distinct clade associated with a robustly supported node (bootstrap support of 100) is formed by two very closely related viruses Wenling crested flounder alphavirus (WCFAV) and Wenling striated frogfish alphavirus (WSFAV), isolated from *Plagiopsetta sp.* and *Antennarius striatus*, respectively.

**Figure 1.**
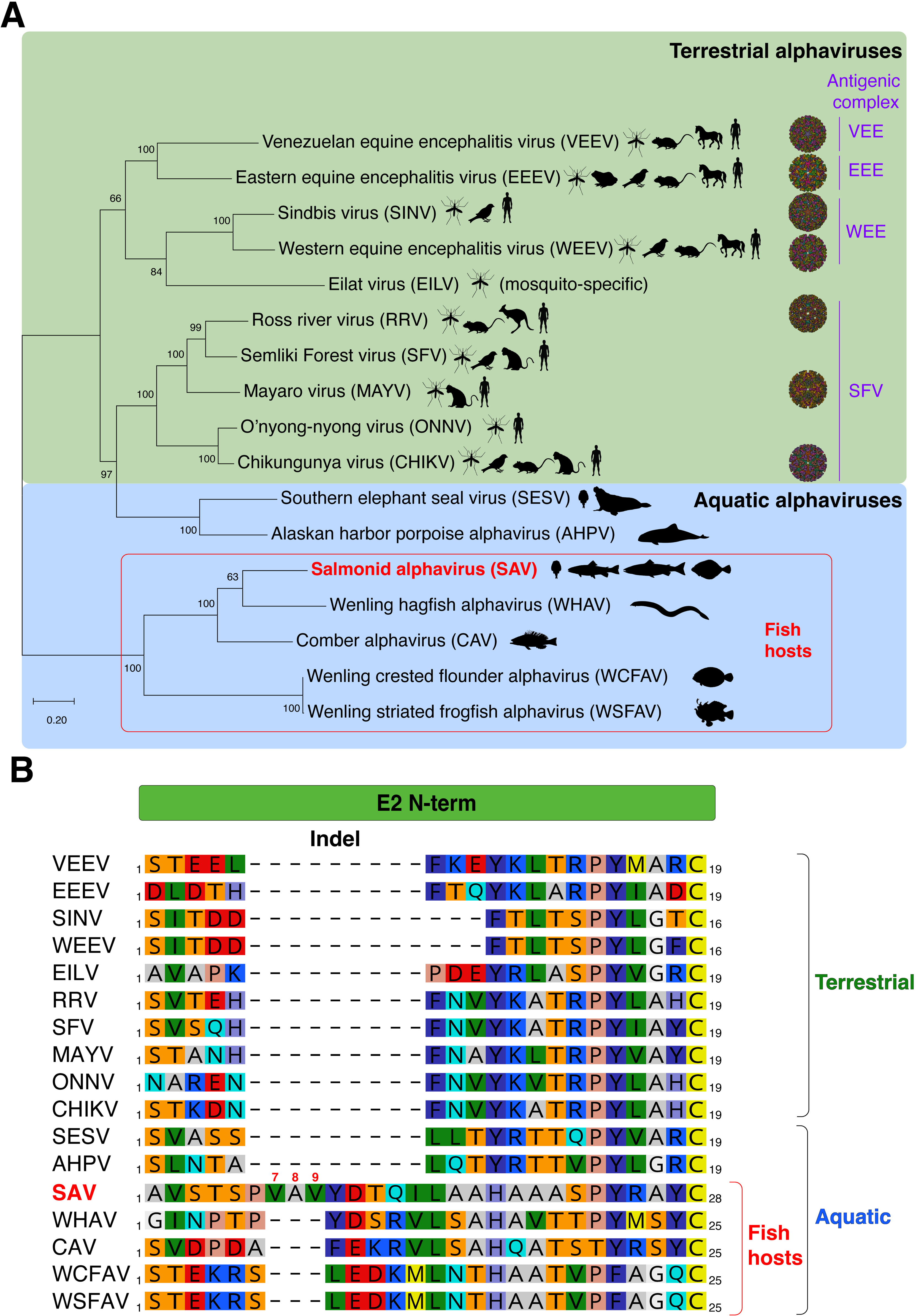
Aquatic and terrestrial alphavirus phylogeny and E2 envelope protein alignment. **A.** Phylogenetic tree of alphavirus structural protein sequences. The amino acid sequence of 17 representative terrestrial and aquatic alphaviruses were aligned and a Maximum-Likelihood (ML) phylogenetic tree was generated (LG (G+I) substitution model). Numbers at nodes indicate percent bootstrap support from 1000 replicates. Tree drawn to scale with branch lengths measured in number of substitutions per site. For each alphavirus, the known host range and vectors are indicated by animal diagrams. Experimentally-determined virion structures are depicted for the following terrestrial alphaviruses: VEEV (PDB <u>3j0c</u>); EEEV (<u>6odf</u>); SINV (<u>6imm</u>); WEEV (<u>8dec</u>); RRV (<u>6vyv</u>); MAYV (<u>7ko8</u>); CHIKV (<u>3j2w</u>). **B.** Amino acid sequence alignment of the N-term region of alphavirus E2. Protein sequences of the 17 aquatic and terrestrial alphaviruses in panel A were aligned and the region corresponding to E2 N-term is shown. The alignment reveals an amino acid insert present only in fish alphavirus E2 proteins (Indel, top). Salmonid alphavirus (SAV) is unique in harboring an additional stretch of 3 residues herein referred to as 7-8-9 triplet based on SAV E2 amino acid numbering.

Several terrestrial alphaviruses, including representatives of the major terrestrial antigenic groupings (VEE, EEE, WEE, and SFV), have been extensively studied structurally. This is attested by the abundance of experimentally-determined whole-virion structures available in the Protein Data Bank (PDB, Figure 1A, right). Importantly, to date there are no experimentally-determined structural data available for alphaviruses infecting fish hosts.

The initial genomic characterization of SAV from rainbow trout (subtype 2), then named Sleeping Disease Virus (SDV), revealed that it diverged substantially compared to its terrestrial counterparts, notably by the length of its individual proteins, in particular the E1 and E2 envelope glycoproteins which contain large inserts ^5^. As displayed in the protein alignment focusing on Alphavirus E2 N-term presented in Figure 1B, a large indel is present close to the N-term with a clear demarcation between alphaviruses infecting fish from the other members of the genus. The alignment highlights the shared insert that is present in fish-infecting alphaviruses. For SAV, the insert is composed of 9 residues with the sequence _6_PVAVYDTQI_14_. Among fish alphaviruses, the SAV insert is unique as it contains an additional stretch of 3 residues _7_VAV_9,_ designated hereafter as triplet 7-8-9. Remarkably, our group has previously demonstrated that a single amino acid substitution at the N-term of SAV2 E2 could dramatically alter virulence *in vivo* in rainbow trout ^11^. The published work reported that a single substitution, Alanine (A) to Valine (V) at position 8 (A8V), was responsible for 90% of the attenuation observed in a recombinant clone previously obtained in the laboratory by reverse genetics. However, upon sequencing verifications performed as part of this current study, we have uncovered that the substitution was erroneously attributed to position 8 (Figure S1). Retrospective RT-PCR and sequencing analyses performed on RNA extracted from preparations of the virulent S49P viral isolate, recombinant SAV (rSAV) strains, as well as plasmid constructs used for SAV reverse genetics all confirm that the attenuating substitution actually concerns position 9 (*i.e.* A9V) and not position 8, with A_9_ associated with virulence and V_9_ associated with attenuation (Figure S1). Of note, the above-mentioned error does not concern the Methionine (M) to Threonine (T) substitution at position 136 (M136T), which was previously found to have a minor role in modulating virulence ^11^. Since position 9 is part of the triplet 7-8-9 triplet within the SAV E2 N-term insert, we decided to investigate this position along with the 7-8-9 triplet further, as detailed below.

### *In silico* structural protein modeling and 3D reconstruction of a SAV virion

To gain more insights into salient structural features of SAV, we set out to model its structural proteins using AlphaFold 3, one of the leading and most accurate deep learning tools for predicting protein structures and complexes ^18^. The polyprotein sequence of the prototype SAV (<u>AJ316246.1</u>) was chosen to predict the structures of E1 and E2 envelope glycoproteins and the capsid (Cp) protein. Since alphavirus E1 and E2 form tight heterodimers and because AlphaFold 3 is able to predict protein complexes, the two proteins were modeled together, while Cp was modeled separately. For Cp, only the structurally-ordered C-terminal (C-term) domain containing a chymotrypsin-like fold was modeled (aa 122-283), since the N-term tail which binds to genomic RNA is disordered ^9,20^. The predicted models of SAV E1, E2, and Cp proteins were then mapped by structural alignment onto a template E2-E1-Cp subunit of VEEV (PDB <u>3j0C</u> ^9^), a reference alphavirus with a robust cryo-EM-determined structure.

AlphaFold 3 generated a model that captures well the architecture of alphavirus E2-E1 heterodimer with similar domain organization (Figure 2A and S2). The overall predicted template modeling (pTM) score, a measure of the accuracy of the entire structure, was high at 0.76 (0-1 range; the higher the better) and the interface predicted template modeling (ipTM) score, which measures the accuracy of the relative position of proteins within a complex, was also high at 0.76 (Table 1). The predicted aligned error (PAE) is an estimate of the error in the pairwise relative position and orientation of protein residues. PAE gives an indication of how accurately AlphaFold positions a given protein subdomain relative to others (Figure S2). For E2-E1 heterodimer, the PAE estimates were generally low except for the TM domains of the proteins. Further, the E2-E1 model was associated with high to very high predicted Local Distance Difference Test scores (pLDDT; 0-100 range, the higher the better) that provide a residue-level confidence metric (Figure S2). The confidence scoring was particularly robust for residues of the beta sheet-rich ectodomains of E1 (aa 1-397) and E2 (aa 1-352) with associated average per-residue Carbon α (Cα) pLDDT (Cα_pLDDT_) scores of 91 and 88, respectively (Table 1). For both E1 and E2, the transmembrane (TM) domains are predicted to be structured as long α-helices with relatively lower average Cα_pLDDT_ scores of 66 to 63, respectively (Table 1).

**Figure 2.**
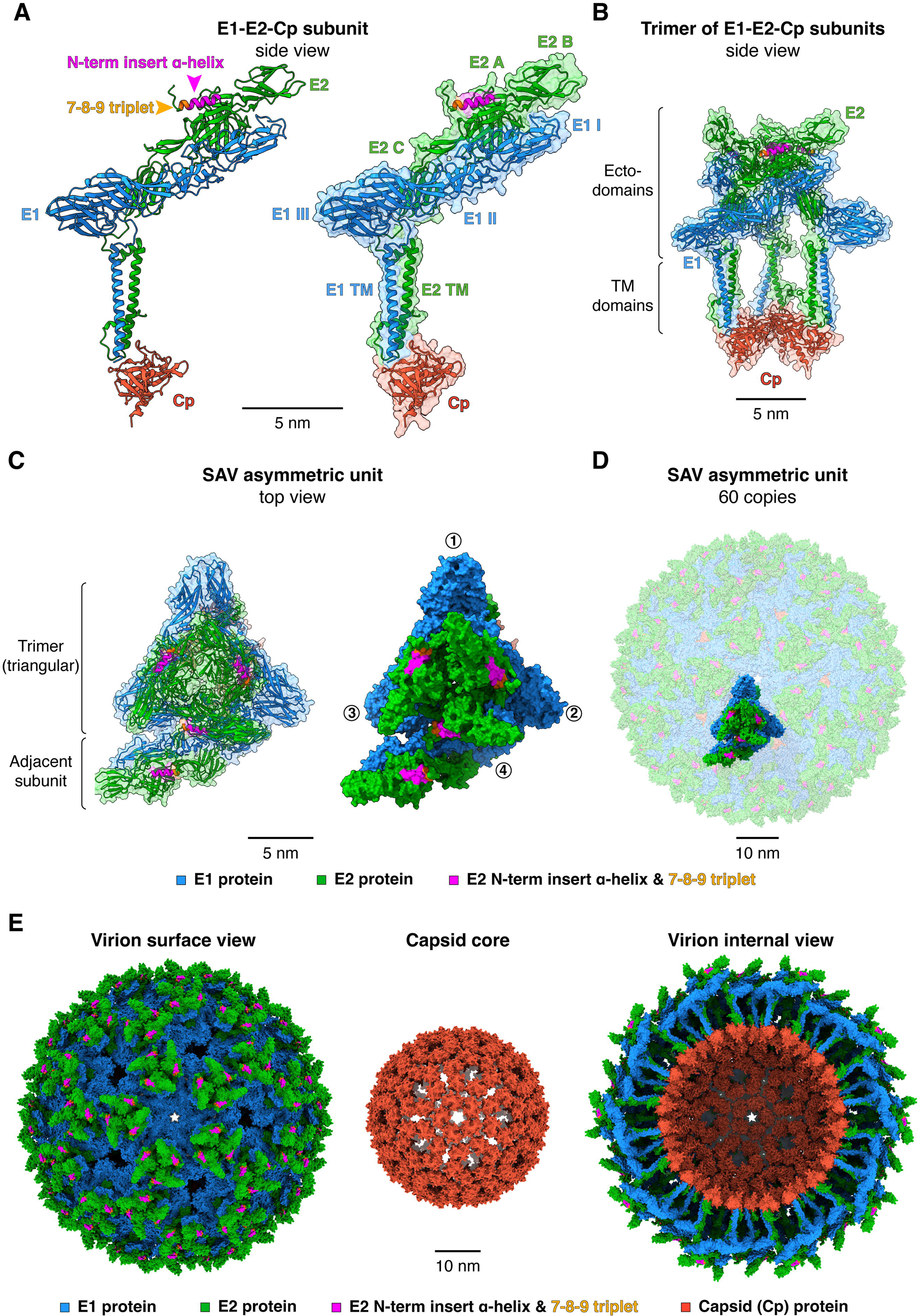
3D reconstruction and architecture of a SAV virion by predictive *in silico* approaches. **A.** E2-E1-Cp subunit shown as ribbons and with surface transparency. The structures of E1, E2 and the ordered C-term half of Cp were predicted using AlphaFold 3. The domains of E1 and E2 proteins are indicated along with the location of E2 N-term insert α-helix and the 7-8-9 triplet (arrows). **B**. Side-view of a trimer of E2-E1-Cp subunits represented as ribbons with surface in transparency. **C**. Details of an SAV asymmetric unit (top view) shown as ribbon and surface representations. Circled numbers indicate individual E2-E1-Cp subunits forming the asymmetric unit. **D**. 60 copies of SAV asymmetric units forming an icosahedral virion. **E**. Predicted structure of a complete icosahedral SAV virus particle with outer surface, capsid core, and internal views. The SAV virion is composed of 240 E2-E1-Cp subunits and assembled using symmetry operations of the reference VEEV icosahedral structure as template (PDB <u>3j0c</u>). For video animations of the virion assembly, see supplementary videos S1-3. Structure labeling, representation, and scale bars were visualized using ChimeraX.

**Table 1.** AlphaFold 3 confidence scorings for SAV E1, E2, and Cp. The table summarizes the different scores generated by AlphaFold 3 for the different regions of the structural proteins of SAV. Scores shown: predicted template modeling, pTM; interface predicted template modeling, ipTM; and the average per-residue Carbon α (Cα) predicted Local Distance Difference Test, Cα_pLDDT_. pTM and ipTM are within the 0 to 1 range (the closer to 1 the better the prediction). Cα_pLDDT_ was calculated by averaging the pLDDT scores of the Cα of each residue within a given protein region (score range 0-100, the closer to 100 the better).

| Protein/complex | Region (aa) | pTM | ipTM | Ca<sub>pLDDT</sub> |
| --- | --- | --- | --- | --- |
| E2-E1 heterodimer | E1 1-461 & E2 1-438 | 0.76 | 0.76 |  |
| E1<sub>ecto</sub> | E1 1-397 |  |  | 91 |
| E2<sub>ecto</sub> | E2 1-352 |  |  | 88 |
| E1<sub>TM</sub> | E1 398-461 |  |  | 66 |
| E1<sub>TM</sub> internal region (AJ316246.1) | E1 429-441 |  |  | 51 |
| E1<sub>TM</sub> internal region (NC_003930.1) | E1 429-440 |  |  | 72 |
| E2<sub>TM</sub> | E2 353-438 |  |  | 63 |
| E2<sub>N-term</sub> α-helix (without E3) | E2 7-20 | | | 63 |
| Cp | Cp 122-283 | 0.87 | - |  |
| Cp | Cp 122-283 |  |  | 90 |
| E3 in uncleaved p62 | E3 1-71 |  |  | 75 |
| E3 | E3 1-71 |  |  | 80 |
| E2<sub>N-term</sub> α-helix without E3 | E2 7-20 | | | 63 |
| E2<sub>N-term</sub> α-helix with E3 and cleaved furin loop | E2 7-20 | | | 85 |
| E2<sub>N-term</sub> α-helix with E3 and uncleaved furin loop | E2 7-20 | | | 86 |

The ordered region of Cp (aa 122-283) was modeled by AlphaFold with high accuracy. It scored a pTM of 0.87, generally low PAE error values, and with a very high average Cα_pLDDT_ score of 90 (Figure 2A, S2 and Table 1).

Intriguingly, the E2 N-term insert and 7-8-9 triplet identified in the protein alignment (Figure 1B) are located in proximity to E2 domain A in a linker region known as the β-ribbon ^8^. Part of the SAV E2 insert and the triplet 7-8-9 are predicted to form an extended α-helix located near the apex of the E2-E1 heterodimer (Figure 2A), with an average Cα_pLDDT_ score of 63 (Table 1). The presence of this α-helix is unexpected as Alphavirus E2 ectodomains are considered to be all-β proteins belonging to the immunoglobulin superfamily, as typified by the CHIKV E2 crystal structure ^8^. To obtain further validation of this atypical α-helical secondary structure, we used RoseTTAFold, another highly accurate protein structure prediction program based on a distinct three-track neural network ^21^, to predict the structure of SAV E2 (Figure S3A). Overall, RoseTTAFold generated a structure of E2 that is highly similar to the one predicted by AlphaFold 3 (Figure S3A). RoseTTAFold also predicts that part of the residues of the N-term insert form an α-helix (E2 aa 9-19). These predictions are further confirmed by PsiPred, a sequence-based secondary structure prediction program which also predicts the presence of an α-helix at E2 N-term (E2 aa 9-20) ^22,23^ (Figure S3B).

Since Alphavirus virions assemble into particles with a regularly-organized architecture adhering to a *T* = 4 icosahedral symmetry ^20^, we reasoned that it could be feasible to reconstruct *in silico* a complete SAV virion from the modeled SAV E1, E2, and Cp structural proteins. To build the SAV virion, the E1, E2, and Cp protein models were first mapped onto a reference alphavirus asymmetric unit (VEEV, PDB <u>3j0c</u>) which is composed of 4 E2-E1-Cp subunits arranged as a triangular trimer (Figure 2B) with an adjacent subunit at its base (Figures 2C). The symmetry operations of the reference VEEV structure allowing to generate a complete icosahedral virion from 60 copies of the asymmetric unit were then applied to the SAV asymmetric unit assembly (Figure 2D). This approach successfully reconstructed an entire SAV virion based on the modeled E1, E2, and Cp proteins. Each modeled proteins fit well within the *T* = 4 reference icosahedral lattice, revealing a typical alphavirus structural organization made of two concentric icosahedral shells with the outer shell composed of the envelope glycoproteins E1 and E2 and an enclosed shell made of Cp proteins (Figure 2E, supplementary videos S1-3). Importantly, the reconstructed SAV virion is composed of 240 copies of each of the structural modeled proteins with the E2 N-term α-helix and 7-8-9 triplet prominently exposed at the surface. Such exposure and repetition suggest that even subtle variations in E2 N-term could potentially lead to measurable phenotypical effects, such as for virulence and neutralizing antibody binding as shown for SAV domain B ^14^. The SAV virion model constitutes a powerful tool to generate testable hypotheses for structure-function studies and for assessing the phenotypical impacts of variations.

### Investigating SAV p62 (E3E2) furin-mediated maturation process

The alphavirus E2 envelope glycoprotein is first synthesized as a precursor called p62 (pE2) which on a sequence level is composed of E2 with a small N-term extension called E3 (71 aa for SAV). E3 plays a pivotal role for p62-E1 heterodimer formation and stabilization in the ER, as its presence avoids premature release of the E1 fusion loop upon exposure to low pH in the compartments of the secretory pathway ^20,24,25^. E3 possesses at its C-term extremity a conserved furin cleavage site (Figure 3A). This basic stretch of residues (consensus sequence R-X-R/K-R) is conserved for all alphaviruses underscoring its functional importance. The maturation of p62 into E2 takes place in the trans-golgi network (TGN) and involves the cellular protease furin which cleaves off E3. In some alphaviruses E3 can remain non-covalently associated with the virion surface, such as for SFV ^26^ and VEEV ^9^. In the case of SAV, because the N-term E2 insert α-helix predicted by AlphaFold 3, RoseTTAFold and PsiPred is located proximal to the C-term of E3 in the protein sequence of p62, we set out to investigate its maturation process and potential consequences for E2 (Figure 3A).

**Figure 3.**
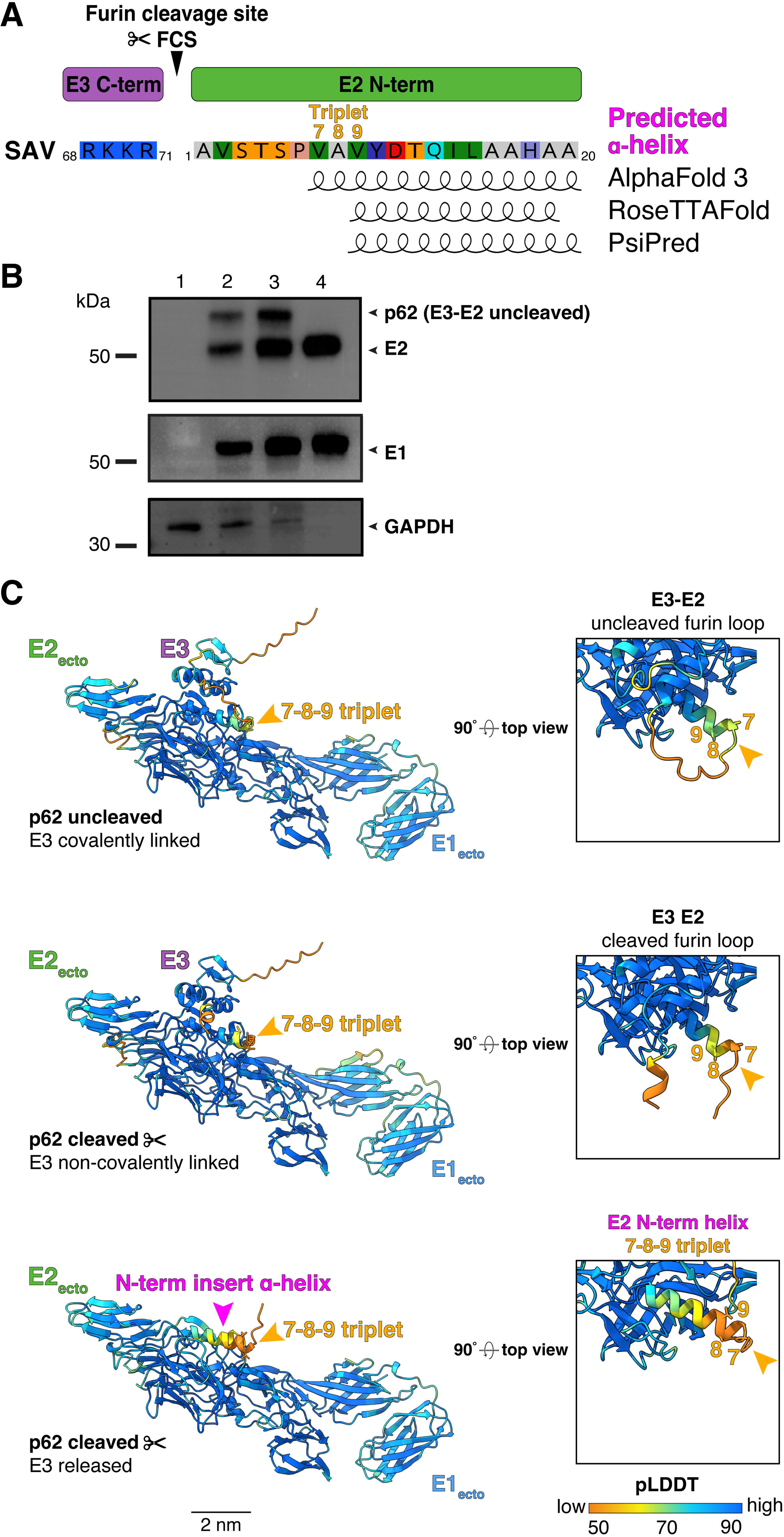
*In vitro* biochemical analysis and *in silico* modeling of SAV E3E2 furin cleavage. **A.** SAV E3 C-term furin cleavage site (FCS) and E2 N-term. Position of the FCS, amino acid composition, triplet 7-8-9 and E2 N-term insert α-helix are shown. The residues involved in the E2 N-term α-helix predicted by AlphaFold3, RoseTTAFold, and PsiPred are depicted as helical cartoons. **B.** Biochemical analysis of cleavage status of p62 in infected cells and purified SAV virions. Western blot analysis was performed on lysates of mock-(lane 1) or SAV2-infected BF-2 cells at an m.o.i. of 1 (lane 2) or 5 (lane 3) or on sucrose cushion-purified SAV2 virions (lane 4). Western blot analyses performed using anti-E2 (17H23) or anti-E1 (78K5) mAbs. GAPDH detection was used as loading control for cell lysates. **C.** Structural modeling of SAV p62 (E3E2) cleavage by furin. SAV E1_ecto_, E2_ecto_, and E3 were modeled as complexes using AlphaFold 3. Top panel corresponds to uncleaved p62 (E3E2), middle panel corresponds to E3 cleaved from E2 after furin cleavage with E3 remaining non-covalently linked, and bottom panel corresponds to E1 and E2 ectodomains after E3 release. Arrows indicate location of E2 N-term ɑ-helix and 7-8-9 triplet. The models are colored using AlphaFold pLDDT confidence scoring.

We first examined biochemically the cleavage status of p62 by performing Western blot analysis of SAV-infected cell lysates and sucrose cushion-purified SAV virions using specific antibodies directed against E1 (mAb 78k5) and E2 (17H23) proteins (Figure 3B) ^14,27^. Both proteins were readily detected in cell lysates and purified virions (Figure 3B, lanes 2-4). The analysis further confirms that in SAV-infected cells where p62 is synthesized and protein maturation occurs, E2 is detected as two distinct bands with one band corresponding to the furin-processed p62 cleaved into E2 and migrating slightly above the 50 kDa marker, and the uncleaved p62 precursor which migrates at a higher molecular weight because of the E3 N-terminal extension (Figure 3B, lanes 2 & 3). In contrast, in purified virions only the ∼50 kDa E2 band is present indicating that only furin-processed E2 and not p62 assembles and buds into newly formed SAV particles (Figure 3B, lane 4).

Next, we computationally simulated three distinct stages of SAV p62 maturation by modeling with AlphaFold 3 the ectodomains of E1 and E2 in presence or absence of E3 (Figure 3C). In a first stage, E3 is modeled as covalently-linked to E2 corresponding to the p62 precursor prior to cleavage by furin at the _68_R-K-K-R_71_`↓`_1_A-V-S-T_4_ cleavage site (Figure 3C, top). In a second stage, E3 is present but non-covalently-linked to E2 corresponding to E3 in complex with E2-E1 ectodomains post furin cleavage (Figure 3C, middle). In a final third stage, E3 is absent and this corresponds to the post-furin cleavage stage and release of E3 from the E2-E1 heterodimer. Remarkably, AlphaFold 3 was able to accurately model E3 with average Cα_pLDDT_ scores of 75 and 80 for the uncleaved (p62) and cleaved form, respectively, with lower scores for the N-and C-termini (Figure 3C, Table 1). SAV E3 is predicted to be structured very similarly to VEEV and CHIKV E3, notably with the presence of two parallel short rod-like α-helices ^8,9^. The predicted models generated by AlphaFold 3 also predict that E3 is positioned near the apex or the E2-E1 heterodimer, on top of the E2 β-ribbon connector where the E2 N-term insert α-helix is located (arrow) and in proximity to E2 domain A, irrespective of whether p62 is cleaved or not (Figure 3C, compare top and middle). The predicted positioning of SAV E3 in the AlphaFold model is reminiscent of the one determined for CHIKV p62 by X-ray crystallography ^8^.

The *in silico* predictive models suggest that for SAV p62, the furin cleavage site forms an exposed, unstructured loop associated with an average Cα_pLDDT_ score below 50, indicative of a disordered and likely flexible region (Figure 3C, top). The furin loop directly connects with SAV E2 N-term and the first residues of the insert α-helix and the 7-8-9 triplet (Figure 3C, top). Remarkably, in all three p62 maturation stages the secondary structure of E2 insert α-helix is predicted to be maintained (Figure 3C). While E3 partially covers the E2 insert α-helix, the triplet residues are exposed during all stages of the maturation process of p62 (Figure 3C top, middle, bottom). Of note, the E2 N-term insert α-helix is predicted with a much higher average Cα_pLDDT_ score of 85 when E3 is modeled together with the E2-E1 heterodimer ectodomains compared to when E3 is absent, average Cα_pLDDT_ score of 63 (Figure 3C middle and bottom, Table 1). This is also the case when the furin loop remains uncleaved (Figure 3 middle with top, Table 1), average Cα_pLDDT_ score of 86. These analyses suggest that the N-term insert α-helix may play an important role in making the furin loop accessible to proteolytic processing and suggests that E3 could also potentially play a stabilizing role locally at the N-term of E2 and the insert α-helix.

## Structural and evolutionary relationships among alphaviruses

To obtain a more comprehensive picture of the structural-evolutionary relationships between aquatic and terrestrial alphaviruses, we examined the E2-E1-Cp subunit of SAV and compared it with that of terrestrial alphaviruses (VEEV and CHIKV) for which experimentally-determined structures are available (PDB <u>3j0c</u> and <u>8fcg</u>, respectively), and AlphaFold 3-predicted models of mammalian aquatic alphaviruses SESV and AHPV, as well as fish-infecting alphaviruses WHAV, CAV, WCFAV, and WSFAV (Figure 4A-D). This comparative analysis reveals that the overall organization of the E2-E1-Cp subunit for all alphaviruses appears to be well conserved. However, a closer inspection of the N-term of the different alphavirus E2 proteins reveals important variations in the N-term insert α-helix that was identified for SAV. Indeed, for the distantly-related terrestrial VEEV and CHIKV, for which cryo-EM structures are displayed, only very short turns are present (Figure 4C). For CHIKV, this short turn was also observed in the X-ray crystal structure of its p62 protein ^8^. For the E2 of aquatic alphaviruses infecting mammals, SESV and AHPV, a short α-helix is predicted to be present (Figure 4C). Similarly to SAV, the closely related fish-infecting alphaviruses, WHAV, CAV, WCFAV and WSFAV, are all predicted to share a longer α-helix at their N-term, similar to the one predicted for SAV (Figure 4D). These results suggest that the N-terminal insert and its predicted extended α-helix is a conserved feature of aquatic alphaviruses which may indicate an important functional and adaptive role for this structural feature for alphaviruses in aquatic environments.

**Figure 4.**
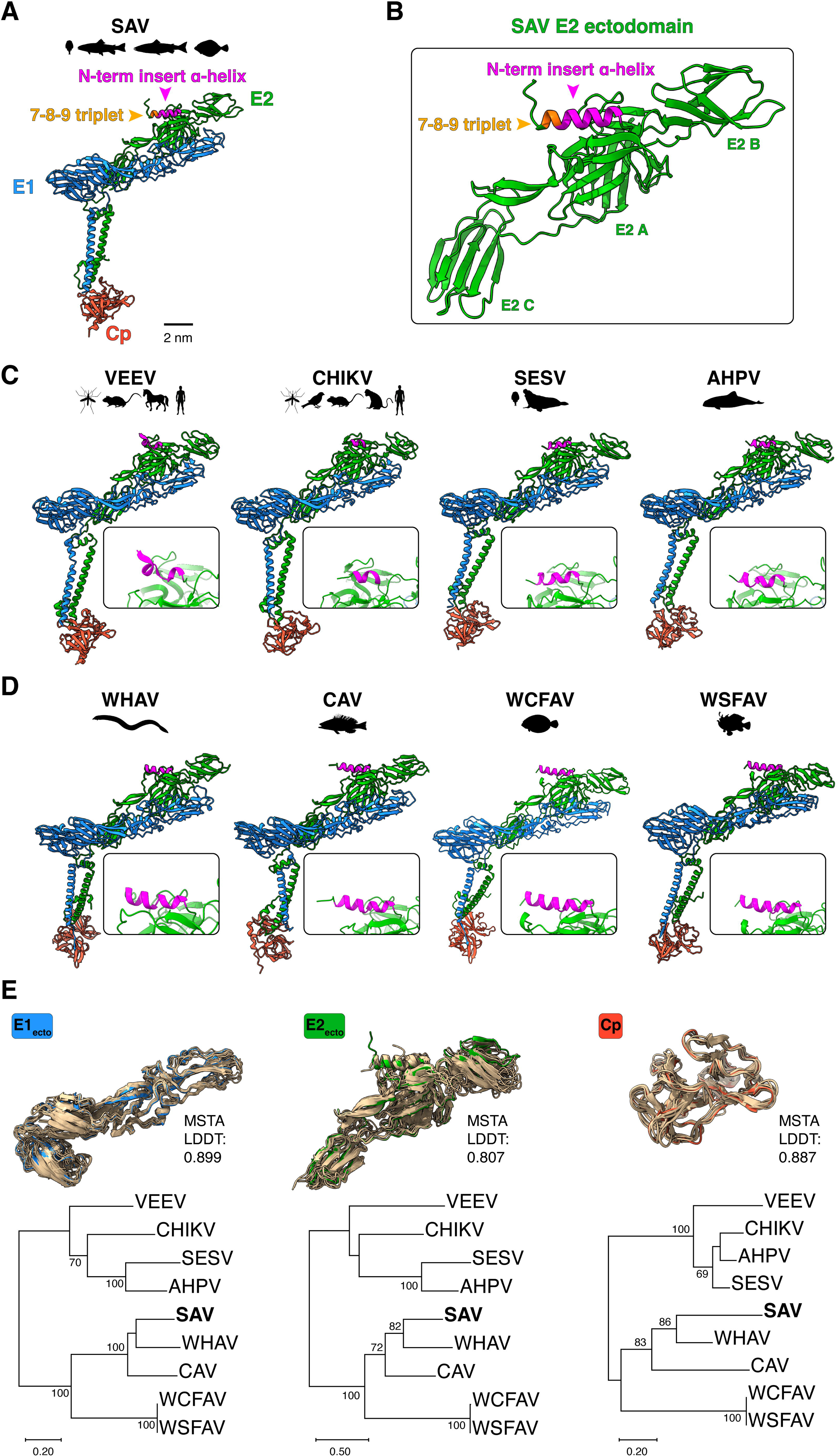
Structural and evolutionary analyses of aquatic and terrestrial alphavirus proteins. **A.** E2-E1-Cp subunit of SAV predicted by AlphaFold 3. **B.** Detailed view of E2 ectodomain with N-term insert helix and 7-8-9 triplet highlighted (arrows). **C.** E2-E1-Cp subunit of VEEV (PDB <u>3j0c</u>) and CHIKV (<u>8fcg</u>) terrestrial alphaviruses and E2-E1-Cp subunit of mammalian aquatic alphaviruses SESV and AHPV predicted by AlphaFold 3. **D.** E2-E1-Cp subunit of WHAV, CAV, WCFAV, and WSFAV fish alphaviruses predicted by AlphaFold 3. For Cp, only the ordered C-term region of the respective proteins was modeled. The known host range and vectors are indicated by animal diagrams. For **C.** and **D.**, a focused view of the E2 N-term of each model is shown in each box. Structure labeling, representation, and scale bar were visualized using ChimeraX. **E.** Comparative analyses between alphavirus proteins using structure-based protein alignment and phylogenetic analysis. FoldMason was used to perform multiple protein structure alignments (MSTA) of alphavirus E1 and E2 ectodomains and Cp protein (C-term domain). In each structural alignment, the blue (E1), green (E2), and red (Cp) protein corresponds to SAV while the superimposed proteins in yellow-gold are from the other species analyzed. The structure-based protein sequence alignments were used to build Maximum-Likelihood phylogenetic trees using WAG+G+I substitution model for E1 and WAG+G for E2 and Cp. Numbers at nodes indicate percent bootstrap support from 1000 replicates. Trees drawn to scale with branch lengths measured in number of substitutions per site.

Next, we assessed the structural relatedness of the ectodomains of E1 and E2, and Cp (structured C-term) of terrestrial and aquatic alphaviruses using FoldMason, a powerful multiple structural alignment algorithm (MSTA) that leverages the structural alphabet developed in FoldSeek ^28,29^. Importantly, FoldMason can handle both experimentally and predicted protein structures. Thus, the E1, E2, and Cp protein domains of terrestrial and aquatic alphaviruses were structurally aligned using FoldMason (Figure 4E top). The structural alignments confirm that protein domains (E1_ecto_, E2_ecto_, Cp) of terrestrial and aquatic alphaviruses all fold very similarly, with few deviations in structural features. For each alignment FoldMason generates a MSTA LDDT score, a measure of similarity of protein structures based on their local structural neighborhoods on a scale ranging from 0 to 1. FoldMason calculated MSTA LDDT scores of 0.899 for E1_ecto_, 0.807 for E2_ecto_, and 0.887 for Cp, indicating that members of the *Alphavirus* genus from aquatic to terrestrial environments share structural similarities for the three analyzed proteins. From the MSTA FoldMason generated a set of 3 structure-informed amino acid alignments. To examine whether these structure-based alignments could recapitulate phylogenetic relationships, each FoldMason structure-based alignment was used to infer a ML phylogenetic tree (Figure 4E, bottom). Overall, the structure-based trees revealed similar branch topologies as sequence alignment-based trees (Figure S4) and recapitulates the main alphavirus groupings obtained in the sequence-based alignment (Figure 1A) with generally high bootstrap support for the major branches (Figure 4E). Of note, for both structure-and sequence-based phylogenetic analyses, the overall tree topologies were similar, but the branches for CHIKV, AHPV and SESV differed slightly in Cp trees compared to E1_ecto_ or E2_ecto_ trees.

### Phylogenetic & structural analysis of SAV E2 and triplet 7-8-9 variations

We next concentrated on naturally-occurring variations and the sequence diversity found at the N-term insert α-helix and triplet 7-8-9 motif. To obtain a sufficiently wide array of SAV isolates from various host species, SAV E2 sequences were retrieved by several methods including performing searches in NCBI Genbank and by conducting Blast analysis using SAV2 E2 reference sequence as input (<u>AJ316246.1</u>). Also included were sequences from SAV field strains made available in published work ^30–32^. This allowed to conduct an extensive phylogenetic analysis of E2 sequences on a diverse set of 90 isolates with representatives from the main subtypes (SAV1-6) from a variety of hosts including rainbow trout, Atlantic salmon, common dab and European plaice (Table S2). As subtype information of some sequences were not provided in the sequence metadata, a nucleotide sequence alignment based on an internal 357-nucleotide E2 fragment used for subtype demarcation was generated ^33^. This allowed to construct a ML phylogenetic tree of 90 SAV isolates (Figure 5A). When available, the host species from which the isolate originated is shown. The E2-based tree allowed to assign subtypes for all 90 isolates. SAV2 and SAV3 isolates, the most represented in this tree, formed a strongly-supported clade (bootstrap support of 99). Likewise, representative isolates from SAV1, SAV4 and SAV5 subtypes clustered into a subclade which was also well supported (bootstrap support of 97) while the SAV6 subtype representative formed a distinct outlier branch from all other subtypes.

**Figure 5.**
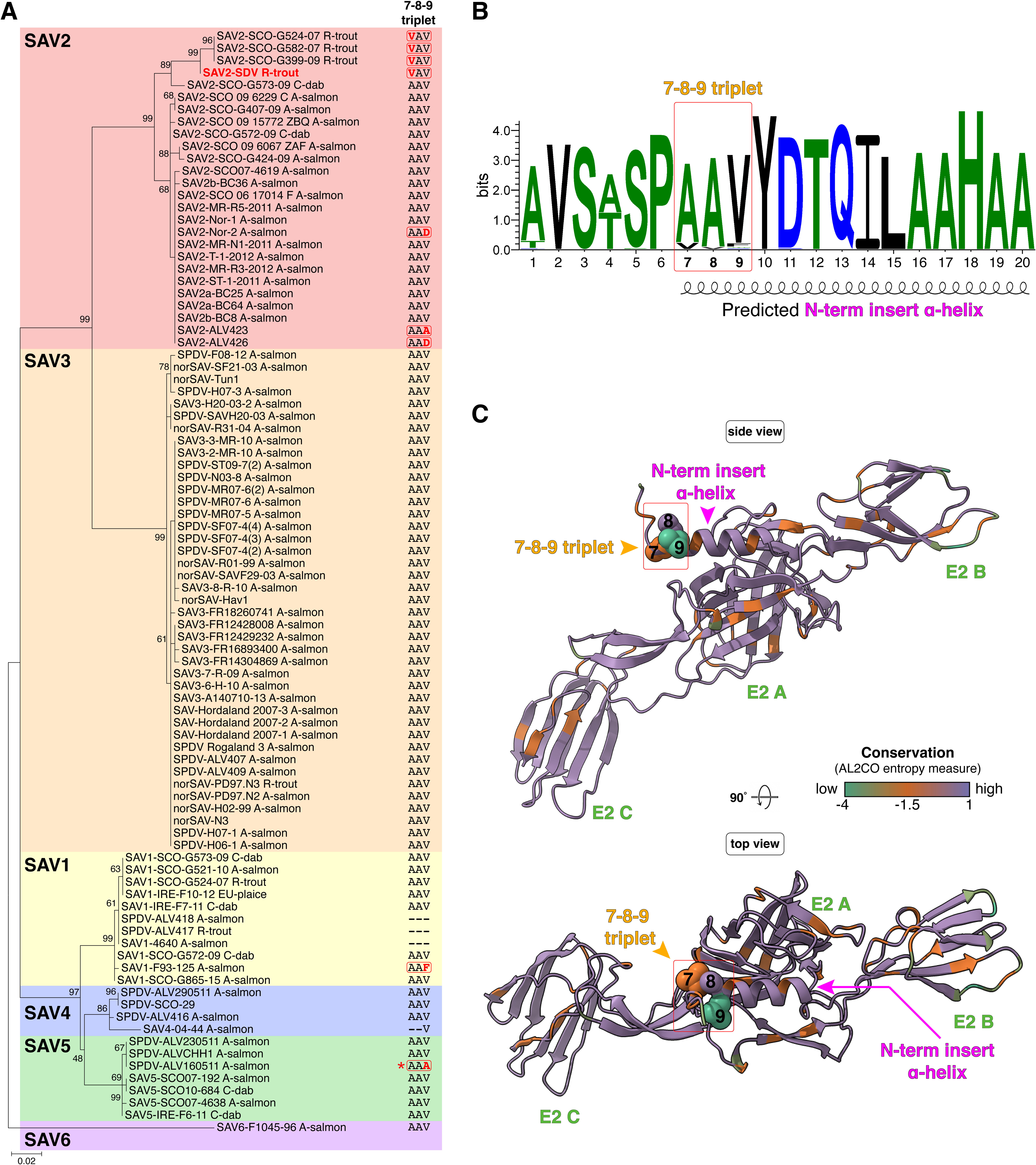
Phylogenetic structural analysis of SAV E2 and triplet 7-8-9 variations. **A.** Phylogenetic tree based on partial E2 nucleotide sequences of 90 SAV strains. SAV E2 sequences (357-nucleotide region used for subtype demarcation) from 90 SAV strains belonging to subtypes 1-6 were retrieved from various sources (Table S2). The sequences were aligned and a phylogenetic tree was generated (GTR (G+I) substitution model). Numbers at nodes indicate percent bootstrap support from 1000 replicates. Tree drawn to scale with branch lengths measured in number of substitutions per site. When known, the host from which the viral sequence originated is indicated. A-salmon: Atlantic salmon (*Salmo salar*); C-dab: common dab (*Limanda limanda*); EU-plaice: European plaice (*Pleuronectes platessa*); R-trout: rainbow trout (*Oncorhynchus mykiss*). For each sequence, the amino acid composition for the 7-8-9 triplet is indicated with variations from the AAV consensus motif highlighted in red/bold. The strain shown in red font corresponds to the SAV reference (<u>AJ316246.1</u>). * denotes the presence of a threonine (T) residue at E2 position 1 for the ALV160511 strain instead of the consensus alanine (A). **B.** Sequence logo of the first 20 amino acids of SAV E2. An alignment of the first 20 amino acids of SAV E2 sequences from the 90 strains shown in panel A was used as input to generate the sequence logo using Weblogo 3.7.12. The total height of a stack at each position indicates sequence conservation, while the height of a symbol indicates the relative frequency of each amino acid. The position of the 7-8-9 triplet is indicated (top). **C.** Mapping of residue conservation within SAV E2 ectodomain. Visualization of residue conservation was performed using the SAV E2 ectodomain (aa 1-352) predicted by AlphaFold 3 and AL2CO entropy measure analysis which is based on a protein alignment of SAV E2 sequences from the 90 strains shown in panel A. Residues within the E2 ectodomain were color-coded according to their conservation with purple indicating a conserved site, orange and green indicating sites associated with increasing variability. The 7-8-9 triplet residues are depicted as spheres. Structure representation and AL2CO analysis were performed using ChimeraX.

For the triplet 7-8-9 site, the most frequently observed motif for all 6 subtypes analyzed was _7_AAV_9_ (A, alanine; V, valine; 77/90), hereafter designated as the consensus motif. The AAV consensus clearly stands out in the Weblogo made by aligning the 90 sequences analyzed in the phylogenetic tree (Figure 5B). Interestingly, the AAV triplet 7-8-9 motif was invariant for all SAV3 isolates. The divergent SAV6 subtype sequence also harbors this conserved sequence. Variations in residue composition were identified for several isolates, notably in 7 isolates of the SAV2 subtype. All variations affected the residues at position 7 or 9, while residue position 8 remained invariant for all isolates analyzed (A_8_). Interestingly, the _7_VAV_9_ motif, containing the Alanine (A) to Valine (V) substitution at position 7 compared to the consensus, was found for SAV2 reference sequence from rainbow trout (<u>AJ316246.1</u>) along with 3 more recently-characterized field isolates from Scottish rainbow trout (SCO-G524-07, SCO-582-07 and SCO-G399-09). Importantly, this specific variation is only found in SAV2 strains infecting rainbow trout, suggesting it may be a host species adaptation within a defined geographic range (Scotland/continental Europe). Interestingly, other variations found for SAV2 strains affect position 9: _7_AAD_9_ (D, aspartic acid) Nor-2 from Atlantic salmon and ALV426 and _7_AAA_9_ for ALV423 strain. Of note, three isolates from subtype 1 SAV, ALV-418 from Atlantic salmon and ALV-417 from rainbow trout as well as strain 4640 from Atlantic salmon presented a 6-residue deletion encompassing the triplet 7-8-9 motif (E2 residues 4 to 9, Δ_4-9_). In addition, the reference SAV1 isolate (F93-125) from Atlantic salmon harbored a variant motif _7_AAF_9_ (F, phenylalanine). All SAV4 strains analyzed harbored the conserved _7_AAV_9_ sequence with the exception of strain 04-44 from Atlantic salmon which had a large 13 amino acid deletion encompassing the E3 furin cleavage site and the E2 N-term up to the first two residues of the 7-8-9 triplet motif. For SAV5 subtype strains only ALV160511 harbored a variant motif _7_AAA_9_. For this strain the motif was associated with another variation at position 1 with an alanine to threonine substitution. This N-term proximal variation is reflected in the Weblogo which reveals that position 4 also displays similar variation (A/T).

Next, sequence conservation was mapped onto the E2 ectodomain structure from the reference sequence (<u>AJ316246.1</u>; _7_VAV_9_ motif) (Figure 5C). The protein sequence alignment of the 90 isolates was used as input to obtain per-residue conservation scores that are mapped onto the SAV E2 ectodomain structure (AL2CO algorithm), an approach that allows to visualize spatial conservation features within a given protein structure (Figure 5C) ^34^. Overall, most of E2 ectodomain amino acids were found to be relatively well conserved (Figure 5C, purple). However, position 9 within the triplet 7-8-9 was associated with a score among the lowest for conservation (green), with position 7 also associated with a low score (orange). This analysis suggests that variations of residue composition within these exposed sites could be of functional and/or structural significance. It is noteworthy to point out that other sites with low conservation are found in unstructured loops of E2 domain B, which is one of the main target sites for neutralizing antibodies (Figure 5C) ^14^.

### *In vitro* investigation of the biological effects of variations in SAV E2 N-term

To assess the biological impact of variations in the structural features identified at the SAV E2 N-term, we used SAV reverse genetics developed in our laboratory to generate recombinant SAV2 (rSAV2) harboring selected mutations identified in the phylogenetic analyses ^35^. In addition to previously obtained triplet 7-8-9 rSAV2 variants which harbor E2 VAV and VAA 7-8-9 triplet motifs (Figure S1) ^11^, other mutations were introduced by site-directed mutagenesis enabling to obtain the following rSAV2 E2 triplet 7-8-9 motif variants: AAV corresponding to the consensus motif, AAA, AAF, and AAD. We also generated the T1-AAA variant which corresponds to the AAA E2 triplet 7-8-9 with a A1T substitution at its N-term, to match the N-term sequence of the ALV16051 strain identified in the phylogenetic analysis (Figure 5A). Of note, despite several attempts, a rSAV2 with a 6-residue deletion encompassing the triplet 7-8-9 motif (rSAV2 E2 Δ_4-9_) was not recoverable (Table S4). rSAV2 harboring the substitutions were recovered and used to infect susceptible BF-2 cells (Table S4). Infection by these rSAV2 were analyzed by immunofluorescence assay at 7-and 10-days post-infection (d.p.i.) to detect SAV-infected cells (Figure 6A).

**Figure 6.**
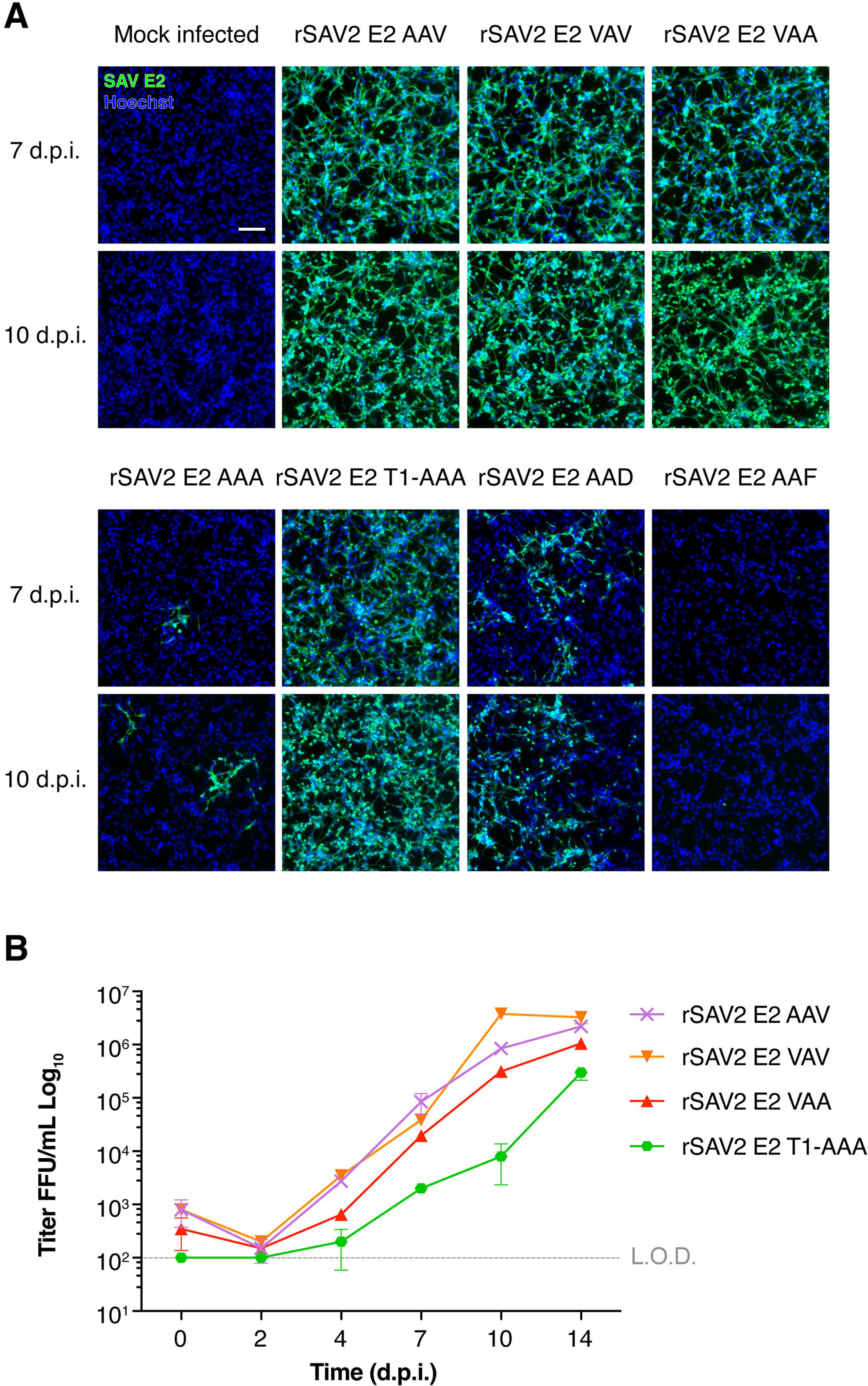
Impact of SAV E2 N-term substitutions on rSAV2 infection in cell culture. **A.** Immunofluorescence assay of BF-2 cells infected with rSAV2 harboring E2 envelope proteins with distinct 7-8-9 triplet motifs: AAV consensus, VAV, VAA, AAF, AAD, AAA, and T1-AAA. E2 N-term variations were introduced using site-directed mutagenesis starting from the VAA variant and recombinant viruses with variant 7-8-9 triplet motif were recovered and used to infect BF-2 cells. Immunofluorescence assays were performed at 7-and 10-d.p.i. Cells were fixed and immunolabeled using anti-SAV E2 mAb (clone 17H23) to visualize infected cells. Scale bar represents 100 µm. **B.** Viral growth kinetics of rSAV2 harboring E2 proteins with variant 7-8-9 triplet motifs. The rSAV2 variants with AAF, AAD, and AAA E2 triplet 7-8-9 motifs were not included in the analysis due to low virus recovery titers. BF-2 cells were infected at an m.o.i. of 0.1. At 0-, 2-, 4-, 7-, 10-and 14-d.p.i. infected cell supernatants were collected, clarified and viral titers were determined by fluorescent focus assay. BF-2 cells were infected with 10-fold serial dilutions of clarified supernatants. After 7 d.p.i. cells were fixed and immunolabeled using anti-SAV E2 mAb (clone 17H23). Fluorescent foci were examined and counted using a fluorescence microscope. Titers are expressed as fluorescent focus units (FFU) per mL. Results are expressed as averages of 2 replicate experiments (n = 2). L.O.D. : limit of detection.

The rSAV2 with the consensus 7-8-9 triplet motif AAV displayed robust infection at 7 and 10 d.p.i. with most cells being positive for infection, a result comparable to infection with the rSAV2 with VAV and VAA triplet motifs (Figure 6A, top). In contrast, infection by rSAV2 with the AAA triplet motif resulted in very limited infection at both time points (Figure 6A, bottom). Strikingly, the introduction of A1T substitution in the triplet AAA background (rSAV2 E2 T1-AAA) restored infectivity to levels similar to those of the consensus sequence (rSAV2 E2 AAV), suggesting a possible compensatory effect. Infection by rSAV2 E2 AAD was associated with poor *in vitro* replication fitness with only patches of infected cells observed at 7 and 10 d.p.i. The rSAV2 E2 AAF variant appeared to be defective and unable to infect cells, with no positively-infected cells observed at both time points (Figure 6, bottom). These immunofluorescence analysis results are confirmed by the titers obtained for each variant (Table S4) and indicate that the triplet 7-8-9 motif is highly sensitive to certain substitutions particularly at position 9 which can have a drastic impact on viral infectivity and replication fitness *in vitro*.

Of the above recovered variant rSAV2, only rSAV2 E2 AAV, VAV, VAA, and T1-AAA yielded sufficient viral titers to conduct viral growth kinetics analyses (Table S4). The replication kinetics measurements were conducted using BF-2 cells infected with the above rSAV2 at a multiplicity of infection (m.o.i.) of 0.1. Supernatants of infected cells were recovered over a period ranging from 0 to 14 d.p.i. and titered by fluorescent focus assay (Figure 6B). This analysis shows that the rSAV2 with E2 AAV consensus triplet motif replicated as efficiently as rSAV2 E2 VAV, reaching similar final titers at 14 d.p.i of 2×10^6^ and 3×10^6^ FFU/mL, respectively. The rSAV2 E2 VAA variant followed a similar growth kinetic and reached a slightly lower final titer of 8×10^5^ FFU/mL. In contrast, the rSAV2 T1-AAA displayed slower replication kinetics, especially at 4, 7, and 10 d.p.i. However, it did reach a final titer of 3×10^5^ FFU/mL at 14 d.p.i. which is close to that obtained for rSAV2 VAA.

### *In vivo* testing of the impact of E2 triplet 7-8-9 in rainbow trout

To examine the impact on virulence of the variations at E2 N-term and 7-8-9 triplet motif *in vivo* during the course of an infection, the rSAV2 analyzed for their replication kinetics were used to infect two distinct lines of juvenile rainbow trout: the INRAE Synthetic (Sy) and AP2 lines (Figure 7). The Sy line was used in previous SAV infection studies allowing for comparative analyses and represents a genetically diverse population while the AP2 line, an isogenic line derived from the Sy line, was chosen because of its high sensitivity to viral infections in particular to hemorrhagic septicemia virus-induced mortality (VHSV) ^36^. For each infection group, 25 rainbow trout were infected with each rSAV2 by intraperitoneal injection (i.p.) at a dose of 10^4^ FFU/fish (average weights: Sy line 2.64 g; AP2 line 2.95 g). For both Sy and AP2 lines, a mock infection control group was included. Mortalities were then recorded over a period of 63 days. For the Sy line infection trial, as expected, infection with the rSAV2 E2 VAA variant, which contains an alanine at position 9 (A_9_) and previously shown to be associated with a virulent phenotype *in vivo* ^11^ (Figure S1) induced mortalities starting at day 25 reaching a cumulative percent mortality (CPM) of 32% at the end of the trial (Figure 7A and Table 2). Infection with rSAV2 E2 VAV variant, with a valine residue at position 9, a substitution previously shown to be attenuating in the Sy line of rainbow trout ^11,35^ (Figure S1), was found to be completely attenuated with no recorded mortalities during the course of the trial. Interestingly, infection by the rSAV2 AAV variant, containing the consensus triplet 7-8-9 motif resulted in 0 recorded mortalities, indicating it is also fully attenuated, like the rSAV2 E2 VAV variant (Figure 7A, Table 2). In contrast, infection with the rSAV2 E2 T1-AAA variant resulted in mortalities, appearing later than infection by rSAV2 E2 VAA (start of mortalities at 38 d.p.i.) but reaching a similar final CPM of 29.2% at the end of the trial. rSAV2 E2 VAA and rSAV2 E2 T1-AAA infections were grouped in the same Log-rank Mantel-Cox statistical group (group A, Table 2) while rSAV2 E2 VAV and rSAV2 E2 AAV infections formed another statistical group (group B, Table 2). These results point to a critical role played by position 9 (alanine to valine substitution) on virulence *in vivo*, in contrast with position 7 where the same substitution has no impact.

**Figure 7.**
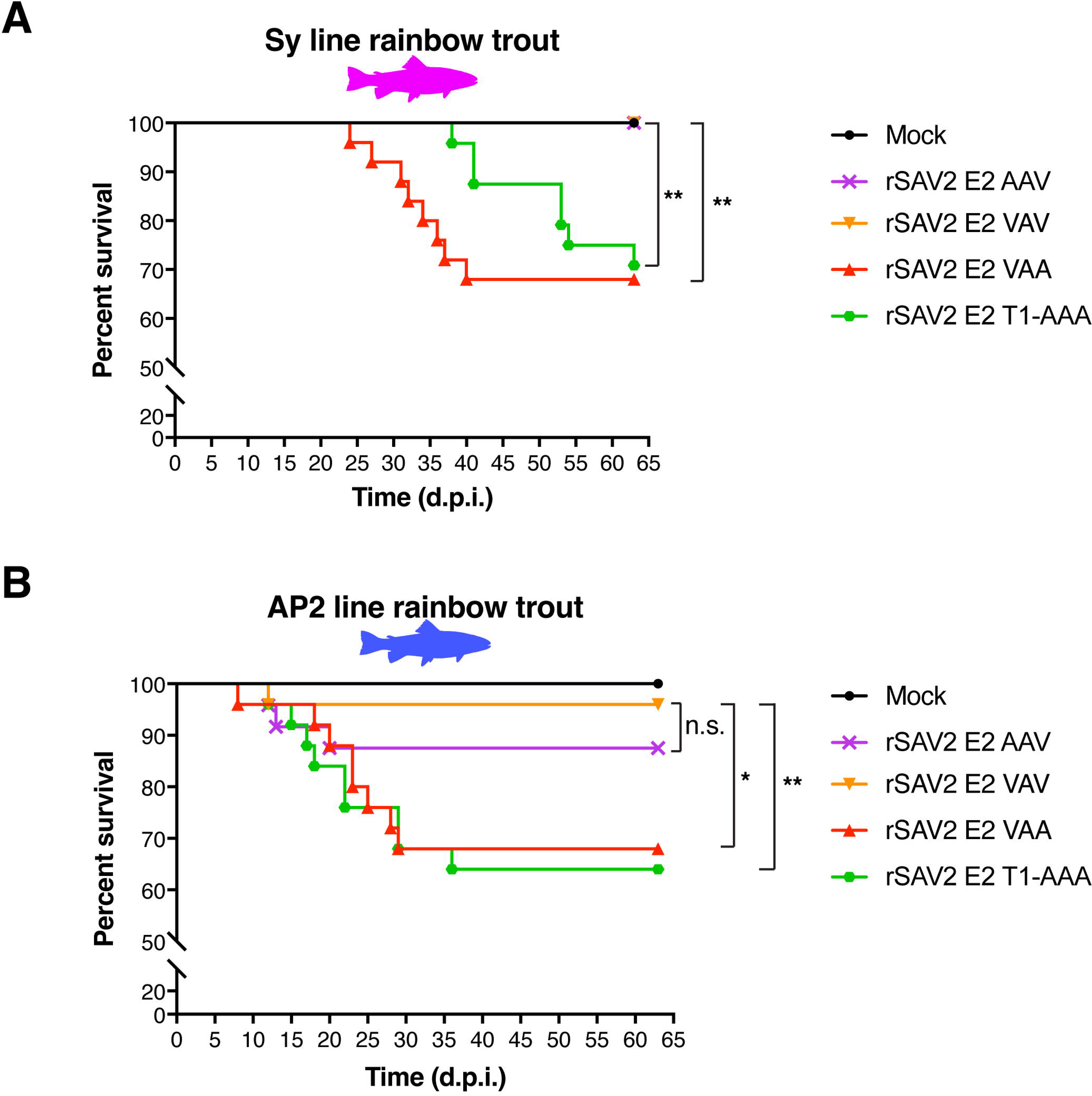
Comparative analysis of the effect of SAV E2 N-term variations in rSAV2 infection in two rainbow trout lines. **A.** Cumulative mortalities induced by rSAV2 infection with variant 7-8-9 triplet motif in the rainbow trout Sy line. Twenty-five juvenile trout (average weight of 2.64 g) were infected by intraperitoneal (i.p.) injection with 10^4^ FFU per fish of the following recombinant viruses: rSAV2 E2 AAV (consensus); rSAV2 E2 VAV; rSAV2 E2 VAA; rSAV2 E2 T1-AAA or were mock infected. Mortalities were recorded daily after injection over a total period of 63 days. **B.** Cumulative mortalities induced by rSAV2 infection with variant 7-8-9 triplet motif in the rainbow trout AP2 line. Twenty-five juvenile trout (average weight of 2.95 g) were infected by i.p. injection with 10^4^ FFU per fish of the same rSAV2 variants as above or were mock infected. Mortalities were recorded daily after injection over a total period of 63 days. For both panels A and B, a comparison of survival between indicated groups was performed using a Log-rank Mantel-Cox test on the Kaplan-Meier survival data. Statistical significance convention used: n.s. *p* > 0.05; * *p* ≤ 0.05; ** *p* ≤ 0.01.

**Table 2.**
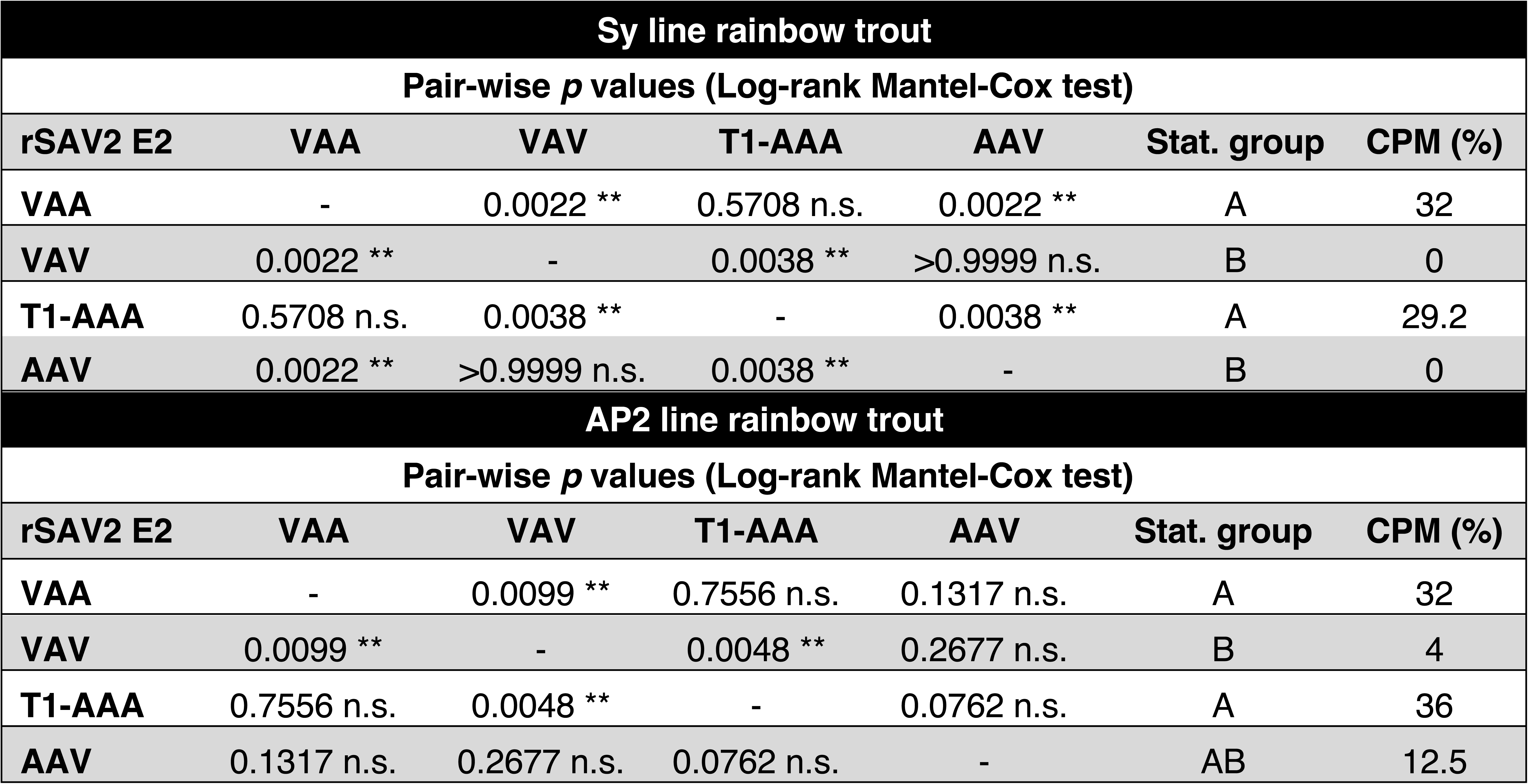
Statistical analyses of rSAV2 infection trials in rainbow trout Sy and AP2 lines. Pair-wise *p* values (Log-rank Mantel-Cox test) calculated from Kaplan-Meier survival curves. CPM, cumulative percent mortality (%); Stat. group: Statistical group. The following convention was used for describing statistical significance: not significant (n.s.) *p* > 0.05; significant (*) *p* ≤ 0.05; highly significant (**), *p* ≤ 0.01.

The infection trial with the AP2 line generally gave similar results as those obtained with the Sy line but with rSAV2 E2 VAA and rSAV2 E2 T1-AAA infections giving rise to mortalities at earlier time points, with a start of mortalities beginning at 8 and 12 d.p.i., respectively (Figure 7B). Both infection with rSAV2 E2 VAA and rSAV2 E2 T1-AAA were associated with similar virulence levels, with 32 and 36% CPM at the end of the trial, forming a single statistical group (group A, Table 2) and (Figure 7B). Moreover, limited mortality was observed for both the attenuated rSAV2 E2 VAV (statistical group B) and rSAV2 E2 AAV infections (statistical group AB) with 4% and 12.5% CPM at the end of the trial, respectively (Figure 7B and Table 2). These results reveal overall higher virulence observed when rSAV2 infect AP2 line compared to the Sy line, a likely consequence of the higher sensitivity of the AP2 line to viral infections as observed in previous infection trials.

To test the genetic stability of recombinant viruses, AP2 rainbow trout that had succumbed post-challenge were sampled to extract viral RNA from their head kidney and spleen. RT-PCR amplification was performed and the E2 N-term region was analyzed by sequencing for positive samples (Figure S5). This confirms that the mutations introduced in rSAV2 E2 AAV, rSAV2 E2 VAA, and rSAV2 E2 T1-AAA remained stable as the respective E2 N-term variations introduced in each recombinant virus were all found to be present with no other mutations found (Figure S5 A-C). In contrast, no positive amplification could be achieved for the only fish that died in the rSAV2 E2 VAV challenge group, putting into question whether the animal succumbed to causes other than rSAV2 infection.

In summary, the *in vivo* infection trials in both Sy and AP2 lines of rainbow trout demonstrate the essential role played by position 9 within the 7-8-9 triplet motif in regulating virulence *via* an alanine to valine substitution, with A9 associated with virulence and V9 associated with attenuation. The same substitution at the proximal position 7 had no impact on virulence.

## Discussion

Compared to their terrestrial counterparts, aquatic alphaviruses have been understudied, with SAV being the only representative species for which a substantial body of research is available, principally due to its disease impact on aquaculture fish species and as a WOAH-listed viral pathogen ^37^. This has left the field with scarce resources to tackle fundamental questions regarding fish alphavirus biology, evolution as well as the mechanisms underpinning host-pathogen interactions, virulence, and interspecies host jumping. Through an integrative approach combining *in silico* 3D reconstruction of an entire SAV virion, phylogenetic and structure-based comparative analyses with *in vitro* and *in vivo* experimental validations, this study lays the groundwork to tackle such questions. As fish alphavirus proteins lack experimentally-determined structures, previous efforts to gain insights were based on classical homology modeling approaches like I-Tasser ^38^ and Swiss-Model ^39^. For example, the former was used to locate a naturally-occurring mutation in SAV3 E2 (P206S) involved in viral fitness and host adaptations ^40^ and the latter enabled to map deletions and residue variations in SAV3 envelope proteins that arise during the course of experimental infection in susceptible hosts ^41^. Similarly, we have previously modeled the non-structural protein 2 (nsP2) of SAV2 using RaptorX ^42^, a template-based protein structure modeling server, in an effort to determine its functional subdomains and investigate its inhibitory function on the RIG-I sensing pathway of the innate antiviral immune response ^43^.

In recent years, the field of structural bioinformatics has seen the rise of revolutionary approaches, such as AlphaFold ^18,19^, RoseTTAFold ^21,44^ and ESMFold ^45^ which are based on artificial intelligence, deep-learning, and protein language models. These advanced programs are able to predict the structures of proteins with high accuracy and at scale. This has led to the establishment of vast databases of predicted structures, such as the AlphaFold Protein Structure Database with over 200 million predicted structures (AFDB, https://alphafold.ebi.ac.uk) and ESM Metagenomic Atlas with over 700 million predicted structures (https://esmatlas.com). However, AFDB excludes viral proteins as it does not handle viral polyproteins like the alphavirus structural and non-structural polyproteins and ESM Metagenomic Atlas lacks taxonomic data severely limiting the study of viral proteins. The Big Fantastic Virus Database (BFVD, https://bfvd.foldseek.com) ^46^ and the Viral AlphaFold Database (VAD, https://data-sharing.atkinson-lab.com/vad/) ^47^ have recently emerged to address this critical gap in predicted viral protein structural data and will greatly benefit advances in virology.

The above tools have already proven their power to study the function and evolution of viral proteins and to shed light on the so-called “viral dark matter” ^48^. This is exemplified in a structure modeling study by Mutz and colleagues on Orthopoxvirus proteins demonstrating multiple cases of exaptation (recruitment) and inactivation of host cellular enzymes by this family of DNA viruses ^49^. In a study on carp edema poxvirus (CEV) for which we took part, a combination of approaches involving structure prediction and structural alignment using the search tool FoldSeek ^29^ was undertaken and revealed the surprising structural relationship between one of its coding sequences, cds46 and host cell endonucleases of fish^50^. This finding hints on a possible cellular origin of cds46 by a gene capture event in a common viral ancestor. The current work underscores the advantage of performing structural alignments, as was done here using FoldMason ^28^ with alphavirus E1, E2, and Cp predicted protein structures to reconstruct structure-based viral phylogenies. Indeed, the rapid evolution of viruses, especially RNA viruses, can be an obstacle to establish clear evolutionary relationships using sequence analysis alone. Because protein structures and folds tend to be more conserved than sequences ^51,52^, structure-based alignments can overcome this limitation and inform on evolutionary relationships over vast time scales. Mifsud and collaborators have recently combined predictive structural bioinformatics with phylogenetic analyses to survey the landscape of *Flaviviridae* family glycoprotein structures ^53^. The authors reveal a complex evolutionary history punctuated by bacterial gene capture and possible recombination events between members of distinct genera.

Thanks to these major advances, the current work takes a holistic approach by modeling entire structures of SAV envelope proteins (E1, E2, and E3), as well as the ordered region of Cp leveraging on the power and accuracy of AlphaFold 3 ^18^. A complete model of an SAV virion was obtained by assembling individual protein models into higher-order hierarchical structures by mapping E2-E1-Cp subunits onto the asymmetric unit and icosahedral lattice of VEEV, a reference alphavirus for which a high-resolution cryo-EM structure has been determined. This constitutes a powerful blueprint for further structure-function studies for SAV and other fish alphaviruses, in particular to examine the functional consequences of mutations and to better understand the biology of aquatic alphaviruses and their evolutionary relationships with terrestrial alphaviruses.

Unexpectedly, modeling of the entire SAV virion revealed that the E2 protein harbored a singular surface-exposed N-term α-helix located in the β-ribbon connecting linker. This helix was confirmed by RoseTTAFold ^21^ and PsiPred ^22^. It stands out because high-resolution X-ray crystallography studies have previously established that CHIKV and SINV E2 ectodomains were structured mostly as all-β proteins belonging to the immunoglobulin superfamily ^8,54^. Further evidence for the N-term α-helix comes from the comparative structural-evolutionary analysis performed in this study showing that the E2 protein of all fish alphaviruses are predicted to contain this feature, which corresponds in part to the shared N-term insert identified in the initial alphavirus E2 protein alignment. Intriguingly, while an N-term α-helix was found to be absent in VEEV and CHIKV terrestrial alphaviruses, a shorter α-helix was predicted for the aquatic alphaviruses infecting sea mammals SESV and AHPV. Since alphaviruses are considered to be of marine origin ^2^, these findings point towards a hypothesis in which the N-term α-helix could represent an ancestral trait conserved in alphaviruses from the aquatic environment. It would be of interest to explore whether the absence of the N-term α-helix in extant representatives of terrestrial alphaviruses was the result of a gradual loss during alphavirus evolution from their original marine environment to their adaptation to terrestrial lifecycle infecting new mammalian, bird, and invertebrate hosts.

The E2 protein alignment highlights that SAV is unique among fish alphaviruses in harboring the 7-8-9 triplet motif corresponding to the first three residues of its E2 N-term α-helix. Previous work conducted in our laboratory using SAV reverse genetics and *in vivo* rainbow trout challenge demonstrated that an A9V substitution in E2 resulted in attenuation (incorrectly assigned to position 8 in ref. ^11^, see Figure S1). This substitution likely arose spontaneously during the establishment of the reverse genetics system for SAV in our laboratory (based on a subtype 2 SAV) ^35^. One of the main challenges towards building a phylogenomic framework is the availability of genomic and metagenomic data that are solidly connected to phylogenetic analyses, epidemiological and experimental data of virulence and fitness ^55^. In a first attempt to apply this framework to SAV, we have undertaken a focused phylogenetic-based approach examining SAV E2 to investigate whether variations found in field isolates and occurring on the triplet 7-8-9 motif of isolates could also play a role in regulating virulence *in vivo*. While clinical metadata such as disease symptoms and severity associated with isolates are still largely lacking, renewed efforts in characterizing SAV genomes have substantially increased the sequence diversity landscape of isolates. This presents an opportunity to bridge the gap between *in silico*, *in vitro* and *in vivo* data on virulence-impacting mutations ^55^.

Examples of how such approaches can lead to insights in the evolution of viral pathogenicity are found in studies on the fish novirhabdovirus infectious hematopoietic necrosis virus (IHNV) that have revealed that cross-species transmission from sockeye salmon to rainbow trout was first accompanied with decreased virulence in its new host followed by a recovery of virulence for some genotypes, which was associated with increased transmissibility ^56,57^. More recently, molecular virulence markers of rainbow trout infection have been characterized scattered throughout the genome of the related viral hemorrhagic septicemia virus (VHSV) ^58^. Much like for SAV, the study showed that single amino acid changes could drastically alter pathogenicity. Phylogenetic analyses also revealed the evolution towards increased virulence for certain subtypes ^59^.

The E2 phylogenetic analysis performed here is based on an extensive set of sequences from 90 SAV isolates. The most frequent triplet 7-8-9 motif observed was _7_AAV_9_ (77/90), which was found in all SAV3 sequences analyzed. For subtypes 1, 2, and 5, variations were observed at positions 7 and 9 but never at position 8 (A_8_) which remained invariant for all isolates analyzed. The E2 protein alignment also revealed that sequences from SAV1 and SAV4 isolates had N-term deletions spanning either a part or the entire triplet motif. Deletions in SAV structural proteins (E1 and E2) appear to be fairly common occurrences as reported by Roh and colleagues in a study based on Nanopore sequencing that followed variations occurring in SAV3 genomes during the course of infection in Atlantic salmon and brown trout ^41^. The majority of the deletions arising spontaneously during infection are believed to be deleterious for the virus. Similar deletion variants were reported by a previous study which was also based on Nanopore sequencing ^60^. These findings echo our inability to recover rSAV2 E2 Δ_4-9_.

Since disease status and severity metadata was lacking for the isolates with triplet 7-8-9 variations, we introduced these variations by site-directed mutagenesis in our rSAV2 system and tested them *in vitro* in susceptible BF-2 cells and *in vivo* in rainbow trout. *In vitro*, the rSAV2 E2 with the _7_AAV_9_ (rSAV2 E2 AAV) triplet, found in the majority of field isolates, had similar growth characteristics as rSAV2 E2 VAA and rSAV2 E2 VAV variants which were obtained previously. However, the capacity to infect BF-2 cell cultures was severely limited for rSAV2 E2 AAA, AAD, AAF variants, possibly due to a loss in viral fitness. Specifically, introduction of a bulky hydrophobic residue (F) or the change to a negatively charged residue (D) at position 9 (AAF and AAD) led to a strong decrease in infectivity. Surprisingly, a more subtle change from AAV to AAA involving alanine and valine small hydrophobic amino acids can also profoundly impact infectivity (AAA), a deleterious effect that is compensated by substituting the small hydrophobic alanine with a polar uncharged threonine at position 1 (T1-AAA).

In contrast to the rSAV2 E2 AAA variant, the rSAV2 E2 T1-AAA was able to infect BF-2 cells albeit with a delay in its growth kinetics compared to rSAV2 E2 AAV. These results suggest possible epistatic effects between triplet 7-8-9 motif residues and the N-term of E2. The recombinants which could replicate efficiently *in vitro* were then tested *in vivo* in two lines of rainbow trout (Sy and AP2). This analysis confirmed that for both rainbow trout lines an alanine to valine substitution at position 9 was accompanied by attenuation and that the same alanine to valine substitution at position 7 had no effects on virulence. From a structure-function vantage point, position 9 can be viewed as a mutationally-sensitive virulence switch. Further testing of other variations at this site either alone or in combination, as found in the different SAV subtypes, could inform whether other residues can alter virulence. As our reverse genetics system is based on the SAV2 subtype, it would also be interesting to perform such analyses using other SAV genetic backgrounds, such as with the rSAV3 backbone.

While our study focused on the triplet 7-8-9 and the first position of E2, it is important to emphasize other sites in E2 as well as other proteins can also impact fitness and virulence. In previous work from our group, another spontaneous E2 substitution, T136M, was shown to have an attenuating effect during *in vivo* rainbow trout challenge, albeit to a lesser degree than the A9V substitution as the latter accounted for 90% of the attenuation ^11^. Close inspection of the protein alignment of the 90 isolates in the current study revealed that none contained a methionine at position 136 (M136) and that T136 is quasi-invariant with only a single isolate, SAV3 8-R10 harboring an alanine, A136 (<u>KC122921.1</u>). However, the impact of the T136A substitution on virulence remains to be elucidated.

During the course of this work, Braaen and colleagues reported to have generated and tested *in vivo* in Atlantic salmon a recombinant virus based on a SAV3 backbone and harboring the A8V and T136M substitutions in its E2 protein ^61^. The authors showed that the recombinant virus was attenuated compared to the parental strain suggesting that an alanine to valine substitution at position 8 could potentially also alter pathogenicity. However, the authors have not tested the contribution of each substitution. Also, the infection models (Atlantic salmon and rainbow trout) and reverse genetics backbones (rSAV3 and rSAV2) are different so the results of Braaen *et al.* may not be directly transposable to this current work. Another case where a single residue change can have a profound phenotypical impact has been studied by Karlsen and collaborators who have identified the P206S substitution in SAV3 E2, which is located in domain B of the protein and appeared during an epizootic episode from a wild reservoir species to farmed salmon in Norway ^40^. The authors tested the substitution using reverse genetics and found that *in vitro* the recombinant virus with the substitution had slower replication kinetics and lower fitness compared to the parental strain but was found to be shed and transmitted more rapidly in the water in an *in vivo* Atlantic salmon cohabitation transmission model.

While we still lack a mechanistic picture for understanding the reasons why substitutions in the N-term of E2 can lead to drastic changes in virulence, a few hypotheses can be put forward. Indeed, the modeling of the entire SAV virion clearly shows that the predicted N-term α-helix containing the triplet 7-8-9 is surface-exposed and repeated 240 times on the icosahedral structure of the virus. As such, it becomes evident that the effects of even a modest modification, such as a single substitution, can be amplified by its repetition across the entire icosahedral surface. In addition, while the receptor of SAV has yet to be identified, the location of these E2 structural features, which are exposed to the external environment, could allow them to be directly involved in host cell receptor interactions.

Furthermore, the close proximity of the triplet 7-8-9 motif to the N-term of E2 suggests it could play a role in interactions with the E3 companion protein, the pH-dependent binding and dissociation of which can impact stability of the E2-E1 heterodimer and exposure of the fusion loop ^62^. During our examination of the different stages of p62 maturation by furin cleavage at the E3E2 interface (_68_RKKR_71_`↓`_1_AVST_4_), we found that the triplet 7-8-9 residues located at the tip of the N-term α-helix were predicted to remain exposed in the presence of E3, whether covalently linked to E2 or not. It is noteworthy to highlight that the furin cleavage site, which is composed of the canonical _68_RKKR_71_ basic motif at the C-term of E3, is predicted to form an unstructured and exposed loop, quite similarly to furin activation loops observed for other viral envelope glycoproteins such as coronavirus spike proteins ^63,64^. To test the effects of substitutions in the E2 N-term region on furin cleavability, we have conducted an *in silico* analysis using ProP 1.0 ^65^ and PiTou 3 ^66^ furin prediction programs (Table S5). This analysis shows that the A1T substitution (T1-AAA variant), which is the closest to the furin cleavage scissile bond (involving residue P1-P1’ of the cleavage site, P1 corresponding to the last residue of E3 and P1’ to the first residue of E2), had a noticeable inhibitory effect on furin cleavability as it scored the lowest prediction values for both ProP and PiTou algorithms. The other mutations involving triplet 7-8-9 residues had little effect on furin cleavage, likely because they are located more distally to the cleavage site.

Regarding the substitution within the triplet 7-8-9 motif, it is remarkable that even subtle changes like A9V can severely impact pathogenicity. Both alanine and valine are uncharged hydrophobic amino acids and have similar biochemical profiles on first approximation. However, they differ in hydropathy scoring, with alanine having the lowest hydropathy score (1.8) among hydrophobic amino acids and valine having the second highest score (4.2), after isoleucine. Furthermore, a notable difference between alanine and valine lies in their side chain structures. While alanine has a straight-chain aliphatic sidechain, valine has a β-branched side chain ^67^. This key difference explains why alanine is considered to be among the best helix-forming residues, while valine is invariably a poor helix former. These distinct properties were used by Gregoret and Sauer to biochemically measure the tolerance of an α-helix of a native protein to combinatorial substitutions with alanine and valine residues. The authors showed that tertiary interactions within the protein mitigate helix destabilization caused by valine substitutions ^67^. Taking into account the biochemical properties of these amino acids, it is reasonable to posit that alanine to valine substitution at position 9 could destabilize the predicted N-term α-helix.

As AlphaFold is unable to predict the effect of single-residue changes on protein structure and stability ^68^, we have performed an additional *in silico* analysis using Pythia, a recently-developed deep learning model capable of rapid and accurate estimation of mutation-associated change in Gibb’s free energy of folding (ΔΔG), a measure of the outcome of an amino acid change on protein stability (Figure S6) ^69^. The AlphaFold 3 predicted structure of the E2 ectodomain (E2_ecto_ aa 1-352, <u>AJ316246.1</u> reference sequence, VAV 7-8-9 triplet motif) was used as input and the program generated a comprehensive heatmap encompassing all residues of E2_ecto_, providing scores at each position and for every amino acid change (Figure S6). A lower score (darker, blue shading) indicates that the substitution is likely to stabilize the protein, while a higher energy value (lighter, yellow shading) indicates that the substitution is more likely to be destabilizing. When focusing on the triplet 7-8-9 motif, mutations on positions 7 and 9 were on the whole more stabilizing (lower scores) while mutations at position 8 were more destabilizing (higher scores). This suggests that position 8 may be less tolerant to variations compared to positions 7 and 9, which is in agreement with the E2 protein alignment and AL2CO scoring showing that position 8 had an invariant alanine, while variations were observed for positions 7 and 9. Regarding the latter positions, Pythia scores for valine to alanine substitutions were negative, -2.3974 and -0.4327 respectively, indicating that these amino acid changes were predicted to be stabilizing. These results are in line with alanine being a helix former and valine as a helix destabilizer. This warrants further biochemical examination of SAV E2 and the N-term α-helix to confirm these results.

This work harnesses the power of structural bioinformatics and lays the foundation for future studies on the evolution of alphaviruses. This is particularly timely as the effects of climate change on sea and freshwater temperatures have already taken by surprise many experts ^70^ and the consequences on the evolutionary trajectories of aquatic viruses remain to be explored. Fish alphaviruses infect ectotherm hosts that have body temperatures that vary according to their environment. Whether these viruses will adapt to increasing temperatures due to climate change is an open question. Our group has already obtained evidence that Novirhabdoviruses, including VHSV and IHNV, can adapt to higher temperatures (unpublished). For fish alphaviruses, indirect evidence that such adaptation is possible can be found in the known temperature ranges of terrestrial alphaviruses such as CHIKV that infect both ectothermic mosquitoes and endothermic mammalian hosts *in vitro* ^71^, indicating a high degree of plasticity in their permissive temperature range. A more direct and tantalizing argument comes from our group’s previous work showing that a variant recombinant virus (rSDV14) was successfully serially passaged in cell culture at 14 °C, well above the temperature of the parental strain originally grown at 10 °C ^35^. Intriguingly, the thermo-adapted strain was shown to gain virulence in a rainbow trout challenge model. The molecular basis for this temperature adaptation and the link with virulence remains to be elucidated. The innovative and integrative approaches used in the current study, notably by mapping virulence-associated mutations on predicted structures combined with *in vitro* and *in vivo* functional testing, can thus be applied to shed light on these phenomena. This would address one of the most challenging threats facing aquaculture globally and, more broadly, would have important ramifications for our understanding of alphavirus ecology and evolution.

## Limitations of the study

A limit of using AlphaFold is that it generates single conformation protein structures. Alphavirus E1 and E2 proteins adopt different conformations during endosomal entry, where low pH triggers conformational changes. Despite this, AlphaFold coherently generated the prefusion conformations of both SAV E1 and E2. These models fitted well with the VEEV icosahedral lattice used for the virion reconstruction and correspond to the virion conformation before cellular entry.

The E2 alignment and phylogenetic analyses introduce an inherent sampling bias due to variations in the number of sequences available depending on the subtype, location, and host. As such, the phylogenetic and sequence variation analyses performed may not closely reflect the frequencies of variations that exist in the field. However, such analysis can provide a qualitative appreciation of substitutions at key positions that can be of functional and evolutionary importance, as highlighted in our study.

We have taken a targeted approach focusing on variations found in SAV E2 N-term. It is important to consider that other sites in E2 and other proteins likely play important roles in modulating virulence, such as E2 A8V, as performed recently by Braaen *et al.*^61^. By leveraging the powerful structural model of SAV we have developed, future studies can extend the functional analyses to other sites for SAV and possibly other related fish alphaviruses.

Here, *in vivo* rSAV2 challenges were performed on rainbow trout, using two distinct lines Sy and AP2, to assess the effects of host genetic background. For future work, it would be of interest to use a salmon model and other rSAV subtypes to gain insights into how E2 N-term variations are linked with host adaptation.

## Resource availability

The 3D reconstructed SAV virion model was deposited in the Model Archive (Swiss Institute of Bioinformatics, SIB; https://www.modelarchive.org) with the following DOI: <u>ma-gkhkz</u>. This deposit also includes the individual models for SAV E2-E1 heterodimer and Cp. The aquatic alphavirus E2-E1-Cp heterotrimer models were deposited with the following DOIs: AHPV, <u>ma-xgzne</u>; SESV, <u>ma-zwpvs</u>; WHAV, <u>ma-</u> <u>md6n0</u>; CAV, <u>ma-zvp1t</u>; WCFAV, <u>ma-6kpi5</u>; and WSFAV, <u>ma-jt78w</u>.

## Acknowledgments

We would like to thank the late Michel Brémont for his guidance and support in the initial stages of this work and acknowledge the members of our team (équipe Virologie Moléculaire des Poissons) for helpful discussions. We gratefully acknowledge Florence Phocas (GABI Unit, INRAE) for proofreading our manuscript. We are very thankful for the valuable advice given by Gerardo Tauriello (Swiss Institute of Bioinformatics) for the SAV virion structure modeling and deposition in the ModelArchive. We warmly acknowledge the support from INRAE’s rodent and fish experimentation unit (IERP, doi.org/10.15454/1.5572427140471238E12), part of the EMERG’IN national research infrastructure (doi.org/10.15454/90CK-Y371), and would like to thank Damien Degallaix, Pénélope Simon, Dimitri Rigaudeau, and Christelle Langevin for their excellent assistance for the *in vivo* studies. Calvin Fauvet is the recipient of a Ph.D. fellowship from the ABIES (Agriculture Food Biology Environment Health) Doctoral School, AgroParisTech. This work was funded in part by young researcher grants awarded to J.K.M. by the Animal Health Department of INRAE.

## Author contributions

Conceptualization: S.B. and J.K.M.

Methodology: S.B., C.F., E.M., J.B., A.L., D.L., and J.K.M.

Investigation: S.B., E.M., and J.K.M

Writing—original draft: S.B. and J.K.M

Writing review and editing: S.B., C.F., E.M., D.L., and J.K.M.

Funding acquisition: J.K.M.

Supervision: S.B. and J.K.M.

## Supplemental information titles and legends

**Figure S1. Verification of SAV E2 N-term sequences from viral preparations and plasmids used for reverse genetics.** Viral RNA was extracted from virus preparations corresponding to **A.** virulent SAV S49P strain (subtype 2), **C.** attenuated rSAV2 previously established in the laboratory, and **E.** virulent recombinant SAV (rSAV-patho). The viral RNA was then retro-transcribed into cDNA using a specific primer targeting E2 then PCR amplified. The samples were then analyzed by Sanger sequencing using specific primers targeting the E2 N-term region. Plasmids used for reverse genetics were also Sanger sequenced using primers that target the E2 N-term region with **B.** corresponding to the plasmid pSAV allowing to recover attenuated rSAV2 while **D.** corresponds to the pSAV-patho plasmid allowing to recover virulent rSAV-patho. **F.** corresponds to the plasmid pJET1.2-*Bsr*GI-patho which contains the E2 N-term mutation responsible for virulence. The sequences and chromatograms were aligned to a reference sequence. Nucleotide and amino acid positions where changes occur are highlighted with color.

**Figure S2. AlphaFold 3 prediction confidence metrics for SAV E2-E1-Cp subunit. A.** Predicted Local Distance Difference Test (pLDDT) scores for SAV E1, E2, and Cp. E1 and E2 proteins were predicted as a heterodimer and Cp was predicted separately (see methods for details). E2-E1 heterodimer and Cp were assembled as a subunit by structural alignment with a reference E2-E1-Cp subunit (VEEV, PDB <u>3j0c</u>). pLDDT corresponds to a local confidence score for each residue position with scores ranging from 0 to 100, the higher the better. **B.** and **C.** Predicted alignment error (PAE) diagrams for E2-E1 heterodimer (**B.**) and Cp (**C.**). PAE is an estimation of the error (expressed Ångströms, Å) in the relative position between two residues and informs on how confident AlphaFold is about the position of a given subdomain relative to others. Lower errors indicated better confidence in prediction. Structure labeling and representation were visualized using ChimeraX.

**Figure S3. SAV E2 N-term structure predictions by RoseTTAFold and PsiPred. A.** SAV E2 protein structure predicted by RoseTTAFold. The protein sequence of SAV E2 (<u>AJ316246.1</u>) was used to generate a protein structure model with RoseTTAFold (https://robetta.bakerlab.org). The protein domains are shown along the E2 protein model. Arrows indicate the location of the predicted N-term α-helix and triplet 7-8-9 motif. **B.** Secondary structure prediction of SAV E2 N-term by PsiPred. The sequence of SAV E2 N-term (<u>AJ316246.1</u>) was used as input to predict secondary structures with PsiPred 4.0 (https://bioinf.cs.ucl.ac.uk/psipred/). The predicted α-helix is shown along with the contributing amino acid residues of SAV E2 N-term.

**Figure S4. Protein sequence-based phylogenetic trees of aquatic and terrestrial E1_ecto_, E2_ecto_, and Cp alphavirus proteins.** The protein sequences of alphavirus E1_ecto_ and E2_ecto_ and Cp (C-term structured domain) proteins were aligned using MAFFT. Each multiple sequence alignments were then used to infer Maximum-Likelihood (ML) phylogenetic trees using WAG+G+I substitution model for E1_ecto_ and WAG+G substitution model for E2_ecto_ and Cp. Numbers at nodes indicate percent bootstrap support from 1000 replicates. Tree drawn to scale with branch lengths measured in number of substitutions per site.

**Figure S5. Analysis of rSAV2 E2 N-term sequences by RT-PCR from viral RNA recovered from infected rainbow trout.** Viral RNA was extracted from the spleen and head kidney of rainbow trout (AP2 line) that had died at different d.p.i. with rSAV2 E2 AAV, rSAV2 E2 VAV, rSAV2 E2 VAA, and rSAV2 E2 T1-AAA. The recovered RNA was retro-transcribed into cDNA and PCR-amplified using specific primers allowing amplification of the E2 N-term region (RT-PCR product size: 473 bp). Positive and specific RT-PCR reactions were obtained for samples of all infected groups except for rSAV2 E2 VAV. For the RT-PCR positive samples **A.** rSAV2 E2 AAV D12, **B.** rSAV2 E2 VAA D18, and **C.** rSAV2 E2 T1-AAA D15, Sanger sequencing was performed using the primers used for cDNA amplification. The days following infection are abbreviated by the letter D followed by the day number. The diagram of SAV E2 N-term features is shown on top of sequencing chromatograms. The sequences and chromatograms were aligned to a reference sequence. Nucleotide and amino acid positions where changes occur are highlighted with color.

**Figure S6. Evaluation of the effects of N-term substitutions on SAV E2 ectodomain stability. A.** SAV E2 ectodomain (<u>AJ316246.1</u>) predicted by AlphaFold 3 with location of the N-term α-helix containing the 7-8-9 triplet. B. Mapping of Pythia scores obtained for SAV E2 ectodomain (https://pythia.wulab.xyz/). Pythia uses protein structures as input and is based on a deep learning model that can predict the energy of each mutation (20 amino acids) at each position within a given query protein. For each position and substitution, Pythia calculates an energy score that provides an estimation of the stability change after such substitution (difference in the Gibbs free energy of unfolding, ΔΔG). Pythia uses a scoring system in which a lower score indicates that the mutation is more likely to be stabilizing, while a higher energy value indicates that the mutation is more likely to be destabilizing.

**Video S1. Video animation of the surface view of the reconstituted SAV virion.** The animation of the surface view of the SAV virion was generated using the record spin movie command in ChimeraX.

**Video S2. Video animation of the capsid core of the reconstituted SAV virion.** The animation of the capsid core of SAV was generated using the record spin movie command in ChimeraX.

**Video S3. Video animation of the internal view of SAV virion.** The animation of the internal view of SAV was generated using the record spin movie command in ChimeraX.

**Table S1. Polyprotein sequences used for *Alphavirus* genus phylogenetic analysis.** The alphavirus polyproteins used to generate the genus-wide phylogenetic tree are shown along with the corresponding NBCI accession numbers and hyperlinks.

**Table S2. Nucleotide sequences used in SAV E2 phylogenetic analysis.** For each SAV sequence, the corresponding SAV subtype, isolate or strain, host species, NCBI accession number or source reference and NCBI raw sequence reads number with hyperlinks are provided. Host species: Rainbow trout (*Oncorhynchus mykiss*), Common dab (*Limanda limanda*), Atlantic salmon (*Salmo salar*), European plaice (*Pleuronectes platessa*).

**Table S3. Primers used in this study.** Shown are the primers used for site-directed mutagenesis, generating a deletion mutant (E2 Δ_4-9_), sequencing, and RT-PCR. For each primer the target region, description, orientation (Forward, F; Reverse, R), and 5’ to 3’ sequences are provided.

**Table S4. Titers of rSAV2 used in this study.** Shown are the rSAV2 with E2 variations and the corresponding titers that were obtained. n.d. not determined because titer is below the limit of detection of 10^2^ FFU/mL. For rSAV2 E2 Δ4-9, despite several attempts the virus was not recoverable.

**Table S5. Effect SAV E2 N-term variations on predicted furin cleavage at the E3E2 junction.** Predicted furin cleavage scores by ProP 1.0 (https://services.healthtech.dtu.dk/services/ProP-1.0/) and PiTou 3 were obtained by using E3E2 junction sequences with E2 N-term variations (E2 position 1 and triplet 7-8-9 motif). ProP scores obtained for sequences above the 0.5 threshold are considered to be predicted to be cleaved. PiTou scores above 0 are predicted to be cleaved (the higher, the better). The furin cleavage motif at E3 C-term is highlighted in bold and blue, while E2 N-term triplet 7-8-9 motif is in bold. Proteolytic cleavage sites are denoted by the following symbol `↓`.

## STAR Methods

### Phylogenetic analyses

Alphavirus and SAV nucleotide and protein sequences were retrieved from NCBI GenBank, Blast searches using SAV2 reference sequence (AJ316246.1), and from published studies (Tables S1, S2 and ^30–32^).

For the terrestrial and aquatic alphavirus phylogenetic analysis, the amino acid sequences of 17 representative alphavirus structural proteins were used (Table S1). The sequences encoding the capsid protein were removed to improve alignment reliability, as this region contains numerous indels and diverges substantially ^2^. The sequences were aligned using MAFFT (v7.388) algorithm ^72,73^ within Geneious software (Biomatters). A part of the protein alignment corresponding to the N-term region of E2 envelope protein was used to highlight conserved and divergent features amongst aquatic and terrestrial alphaviruses, notably an insert only present in fish alphavirus and the unique 7-8-9 triplet motif of SAV. Phylogenetic analyses were performed in MEGAX 10.1.8 ^74,75^. The choice of the evolutionary model was determined from 56 models based on the Bayesian information criterion (BIC) and a Maximum-Likelihood (ML) tree was generated from the alignment using LG (G+I) substitution model ^76^. Phylogeny testing was undertaken by the bootstrap method with 1000 replicates. The phylogenetic tree was drawn to scale in MEGAX and formatted using FigTree 1.4.4 with branch lengths measured in the number of substitutions per site. Numbers at nodes indicate bootstrap support.

For the SAV phylogenetic analysis, nucleotide sequences corresponding to an internal 357-bp region of E2 used for SAV subtype demarcation were retrieved from 90 representative SAV isolates (Table S2) ^33^. When available, the host species from which the viral sequence was sampled was noted and included in the tree. All major SAV subtypes (1-6) are represented in the analysis. Verifications performed on a SAV7 sequence (isolate F/58/17 ^77^) showed that it contained numerous nonsense mutations rendering the encoding proteins non-functional and was not included in the analysis. The nucleotide sequences were aligned (codon-aware) in MEGAX using MUSCLE ^78^. A Maximum-Likelihood (ML) tree was generated from the alignment using the GTR (G+I) substitution model. Phylogeny testing was performed using the bootstrap method with 1000 replicates. The phylogenetic tree was drawn to scale in MEGAX and formatted using FigTree 1.4.4 with branch lengths measured in the number of substitutions per site. Numbers at nodes indicate bootstrap support.

A protein sequence alignment of the E2-encoding region of the above 90 SAV isolates was generated in Geneious Prime 2025.0.3 (Biomatters) using MAFFT (v7.388) algorithm. This protein alignment was used for subsequent analyses, including determining the amino acid composition of the 7-8-9 triplet for each isolate analyzed in the SAV phylogenetic tree, the sequence logo analysis (WebLogo) of E2 N-term, and mapping of conservation on predicted E2 structure (AL2CO entropy measure) (see below).

### AlphaFold-based fish alphavirus protein structure predictions

The predicted structures of fish alphavirus E1, E2, and capsid (Cp) proteins were modeled using AlphaFold 3 (https://alphafoldserver.com) ^18,19^. For capsids, only the C-term portion of the proteins were modeled to match with the structurally ordered domain of experimentally-determined alphavirus structures (*e.g.* VEEV PDB <u>3j0c</u>) ^9^. For SAV, the predictions were based on the reference SAV2 protein sequences (<u>AJ316246.1</u>). Since alphavirus E1 and E2 envelope proteins form a closely-associated heterodimeric complex and as AlphaFold 3 can natively generate multimeric models, SAV E1 and E2 proteins were modeled together as a heterodimer, while the C-term half of the capsid (aa 122-283) was modeled separately. The overall prediction scores for both the E2-E1 heterodimer and capsid were high with pTM scores of 0.76 and 0.87 respectively (Table 1). However, for E1, AlphaFold 3 gave poor pLDDT scores for a stretch of residues in the alpha helical transmembrane (TM) domain (aa _429_RIVGNPSGPVSSS_441_, average Cα_pLDDT_ of 51, Table 1) introducing a kink and break in the alpha-helix which are not present in any of the experimentally-determined structures of alphavirus E1 proteins (*e.g.* VEEV PDB <u>3j0c</u> and CHIKV PDB <u>8fcg</u>), indicative of a relatively high degree of prediction error. To improve the predicted model of SAV E1, this stretch of amino acids was replaced with that of the SAV2 E1 sequence that was previously cloned in our laboratory (aa _429_GIVGTLVVLFLI_440_) and found in other SAV strains (*e.g.* <u>NC_003930.1</u>) ^35^. By doing so, AlphaFold 3 was able to predict a TM domain with a linear alpha-helix with improved pLDDT values (average Cα_pLDDT_ of 72 for the internal TM region, Table 1) and matching experimentally-determined alphavirus E1 structures (Figure S2). This improved predicted model of SAV E1 was subsequently used throughout this study.

To confirm the predicted N-term α-helix predicted by AlphaFold 3 in SAV E2, the protein sequence of SAV E2 was used to generate a full-length protein model by RoseTTAFold, another leading structure-prediction algorithm (https://robetta.bakerlab.org) ^21^. In addition, the N-term sequence of E2 was analyzed using PsiPred (https://bioinf.cs.ucl.ac.uk/psipred/) ^79^, a PSI-BLAST-based secondary structure prediction algorithm to predict secondary structures.

For the analysis of E2-E3 furin cleavage, the E3 sequence (<u>AJ316246.1</u>) was modeled using AlphaFold 3 together with E2 and the above-mentioned E1 (with modified TM domain). Modeling of E3 was performed either as the uncleaved form (p62, E3E2), or as the cleaved form with E3 non-covalently associated with E2.

For the other aquatic alphaviruses infecting mammalian or fish hosts, the NCBI source accession numbers of protein sequences used for modeling the E2-E1 heterodimer and capsid (C-term structurally ordered portion) were the following: Southern elephant seal virus (SESV) <u>AEJ36233.1</u> sampled from *Mirounga leonine*; Alaskan harbor porpoise alphavirus (AHPV) <u>QJE50388.1</u>, sampled from *Phocoena phocoena*; Comber alphavirus (CAV) <u>MN207265.1</u>, sampled from *Serranus cabrilla*; Wenling crested flounder alphavirus (WCFAV) <u>MG600127.1</u>, sampled from *Plagiopsetta sp.*; Wenling hagfish alphavirus (WHAV) <u>MG600128.1</u>, sampled from *Eptatretus burgeri*; and Wenling striated frogfish alphavirus (WSFAV) <u>MG600126.1</u>, sampled from *Antennarius striatus*. To generate structural models with AlphaFold 3, E1, E2 and Cp were modeled as a heterotrimer. For capsids, the amino acid boundaries used were based on protein alignments and boundaries used for SAV Cp: SESV aa 108-268; AHPV aa 76-236; CAV aa 142-303; WCFAV aa 136-296; WHAV aa 167-327; and WSFAV aa 136-296. The overall pTM scores obtained for the AlphaFold 3-predicted heterotrimeric complexes were the following: SESV 0.74; AHPV 0.72; CAV 0.68; WCFAV 0.69; WHAV 0.66; and WSFAV 0.68.

The experimentally-determined structures of VEEV and CHIKV were retrieved from PDB (<u>3j0c</u> and <u>8fcg</u>). Molecular visualizations were performed using UCSF ChimeraX 1.9 ^80^.

### *In silico* reconstruction of a whole SAV virion

To reconstruct a 3D model of an entire SAV virion, the predicted structures for E1, E2, and the capsid proteins generated by AlphaFold 3 were mapped onto the icosahedral lattice of Venezuelan equine encephalitis virus (VEEV), a reference alphavirus structure which has been experimentally-determined by cryo-EM (PDB <u>3j0c</u>, *T* = 4 icosahedral symmetry) ^9^. The TM domains of SAV E1 and E2 envelope proteins, which are connected to the ectodomains by a flexible linker, were structurally aligned to obtain a better alignment fit with the corresponding regions of VEEV E1 and E2 proteins using ChimeraX ^80,81^. Further refinements to detect and avoid clashes within and between SAV E2-E1-Cp subunits were performed using Isolde ^82^. The models of SAV E1, E2, and Cp were then fitted by structural alignment (MatchMaker tool within ChimeraX ^80,81^) onto the asymmetric unit of VEEV virion, which is composed of 4 copies of E2-E1-Cp heterotrimer (PDB <u>3j0c</u>). To apply the symmetry operations generating a complete icosahedral virion from a single SAV asymmetric subunit (240 copies of E1, E2, and Cp proteins), the following list categories of the VEEV PDB <u>3j0c</u> structure file were copied in the SAV asymmetric unit mmCIF file: _pdbx_struct_assembly, _pdbx_struct_assembly_gen and _pdbx_struct_oper. All molecular visualizations were performed using UCSF ChimeraX 1.9 ^80^.

### Structure-based phylogenetic alignment and trees

FoldMason was used to generate structure-informed phylogenetic trees, using the AlphaFold 3 predicted structures of the ectodomains of E1 and E2 and the C-term portion of Cp of aquatic alphaviruses infecting mammals or fish hosts (https://search.foldseek.com/foldmason) ^28^. FoldMason is a fast and accurate multiple structure aligner (MSTA) based on Foldseek’s 3Di alphabet ^29^. For each of the predicted protein structures used as input, the following aa boundaries (defined by protein sequence alignments with terrestrial alphaviruses) were used for improved structural alignment: SAV E1_ecto_ aa 1-397, E2_ecto_ aa 1-352, Cp_C-term_ aa 134-283; SESV E1_ecto_ aa 1-377, E2 _ecto_ aa 1-334, Cp_C-term_ aa 121-268; AHPV E1_ecto_ aa 1-377, E2_ecto_ aa 1-336, Cp_C-term_ aa 89-236; CAV E1_ecto_ aa 1-397, E2_ecto_ aa 1-352, Cp_C-term_ aa 155-303; WCFAV E1_ecto_ aa 1-396, E2_ecto_ aa 1-354, Cp_C-term_ aa 148-296; WHAV E1_ecto_ aa 1-397, E2_ecto_ aa 1-350, Cp_C-term_ aa 179-327; and WSFAV E1_ecto_ aa 1-396, E2_ecto_ aa 1-354, Cp_C-term_ aa 148-296. For the terrestrial alphaviruses VEEV and CHIKV (PDB <u>3j0c</u> and <u>8fcg</u>), the following aa boundaries were used: VEEV E1_ecto_ aa 1-380, E2_ecto_ aa 1-342, Cp_C-term_ aa 127-275; and CHIKV E1_ecto_ aa 1-379, E2_ecto_ aa 5-342, Cp_C-term_ aa 111-261. FoldMason generates a guide tree, a graphical representation of the structural alignment, as well as a structure-based alignment (MSTA). For each protein (E1_ecto_, E2_ecto_, and Cp_C-term_), the MSTA was used to generate a Maximum-Likelihood (ML) phylogenetic tree. Phylogenetic analyses were performed in MEGAX 10.1.8 ^74,75^. For each protein, the choice of the evolutionary model was determined from 56 models based on the Bayesian information criterion (BIC) and a Maximum-Likelihood (ML) tree was generated from the alignment using WAG+G+I substitution model for the E1_ecto_ tree and WAG+G substitution model for the E2_ecto_ and Cp_C-term_ trees ^76^. Phylogeny testing was undertaken by the bootstrap method with 1000 replicates. The phylogenetic tree was drawn to scale in MEGAX and formatted using FigTree 1.4.4 with branch lengths measured in the number of substitutions per site. Numbers at nodes indicate bootstrap support. Equivalent sequence-based trees were generated for comparison (see Figure S4).

### Visualization of SAV E2 sequence conservation

To obtain a representation of the sequence conservation at the N-term of SAV E2, a sequence logo of the protein was generated using Weblogo 3.7.12 (https://weblogo.threeplusone.com/create.cgi) ^83^. A protein alignment focused on the first 20 amino acids of SAV E2 was derived from the full-length protein alignment of the 90 SAV E2 sequences previously generated (see above). This alignment was used as input for the Weblogo server. The total height of stacked symbols at each position indicates the degree of conservation at that position and the height of each symbol within a given position indicates the relative frequency of each amino acid.

To map conservation onto the structure of the SAV E2 ectodomain (aa 1-352) predicted by AlphaFold 3, the ALCO2 entropy measure program was implemented in ChimeraX 1.9 ^34,80^. Based on the previously obtained protein alignment of 90 SAV E2 sequences (see above), AL2CO calculates a conservation index for each residue of the alignment, which can be mapped and color-coded onto each position of SAV E2 ectodomain ^34^.

### Predicted impact of N-term variations on SAV E2 protein stability

Calculation of the predicted effect of substitutions on protein stability was performed using Pythia (https://pythia.wulab.xyz/), a robust deep-learning model able to rapidly and accurately predict the effect of mutations on protein stability using protein structural information as input ^69^. The input structure used is the E2 ectodomain (aa 1-352) of the reference SAV (<u>AJ316246.1</u>, with A_1_ and _7_VAV_9_ triplet 7-8-9 motif). Pythia generates a heatmap and numeric table covering all positions of a given protein and tests at each position the effect of substitutions with each of the 20 amino acids. For each position and residue substitution, Pythia attributes a score proportional to ΔΔG, which corresponds to the difference in the Gibbs free energy of unfolding (ΔG) between two proteins (non-substituted *vs.* substituted). A lower score indicates that the substitution is more likely to stabilize the protein, while a higher score indicates that the substitution is more likely to destabilize the protein.

### Plasmid constructs and site-directed mutagenesis

The plasmids pSAV-patho (pathogenic, originally named pSDVStruct_wt_ ^11^) and pJET1.2-*Bsr*GI_patho were generated in our laboratory and were used here to recover recombinant SAV (rSAV) with variant E2 N-term amino acid composition ^11,14,35^. pSAV-patho is an infectious plasmid construct containing full-length SAV cDNA (subtype 2) and the plasmid pJET1.2-*Bsr*GI_patho consists of a *Bsr*GI-digested DNA fragment encompassing the complete E2 coding sequence and cloned into the pJET1.2 vector. Both pSAV-patho and pJET1.2-*Bsr*GI_patho contain the E2 substitutions found in the virulent strain of SAV, SDV S49P (Figure S1). Sanger sequencing (Eurofins Scientific) with SEQ13 and pJET1.2For primers, respectively, demonstrates that pSAV-patho and pJET1.2-*Bsr*GI_patho plasmids contain an E2 sequence with VAA triplet 7-8-9 (Figure S1 and Table S3). The first reverse genetics plasmid pSAV (originally pSDV) contains the VAV triplet 7-8-9 (Figure S1). Targeted mutagenesis of SAV E2 N-term was accomplished using QuickChange Site-Directed Mutagenesis Kit (Agilent Technologies) according to the manufacturer’s guidelines with the pJET1.2-*Bsr*GI_patho as template and the mutagenesis primers SDM64-68 (Table S3) to generate the following E2 variants: AAV, AAF, AAD, AAA and T1-AAA. To generate an internal 6-residue E2 N-term deletion mutant (E2 Δ_4-9_), a PCR amplification-based strategy using a 5’-phosphorylated internal primer pair (Table S3) was used as described by Sourimant *et al.* ^84^. Targeted mutations and the deletion were verified by Sanger sequencing (Eurofins Scientific) using the sequencing primer SEQ13 (Table S3). Each of the mutated *Bsr*GI DNA fragments were cloned back into the *Bsr*GI-digested pSAV-patho backbone.

### Cell culture and recombinant SAV (rSAV) recovery

BF-2 cells derived from Bluegill fry (*Lepomis macrochirus*) were maintained at 14 °C in Glasgow Minimum Essential Medium (GMEM, Pan Biotech) supplemented with 23.75 mM Tris-HCl pH 7.6, 4 mM NaHCO_3_, 2 mM L-glutamine (Eurobio), 10% fetal bovine serum (Capricorn Scientific), 1% tryptose phosphate broth (Gibco ThermoFisher) and penicillin-streptomycin antibiotics, respectively at 100 U/mL and 100 µg/mL (Biovalley).

For recovery of rSAV2 with variant E2 N-term, BF-2 cells were trypsinized (Difco Trypsin, BD Biosciences), counted and seeded at density of 5×10^6^ cells/well in six-well plates and transfected by electroporation (Amaxa, Lonza) with 2 µg of each of the mutated pSAV constructs to generate rSAV2 E2 AAV, AAF, AAD, AAA, and T1-AAA variants using the T solution and the T-020 electroporation program. Cells were incubated at 20 °C overnight in GMEM medium supplemented with 10% FBS. 24 h post-transfection, the supernatants were replaced with serum-free GMEM medium and the cells were incubated at 10 °C for 6–10 days. rSAVs with variant E2 N-term were amplified through two successive passages on BF-2 cells seeded in 24-well plates. Viruses were titrated by immunofluorescence assays as described below.

Verification of the E2 N-term sequences of SAV (SDV) S49P, rSAV2 and rSAV-patho (previously named rSAVStruct_wt_ ^11^) were carried out by performing viral RNA extractions from cell culture supernatants using QIAamp kit (Qiagen). Viral RNAs were retrotranscribed using SuperScript reverse transcriptase (Invitrogen) and the *Bsr*GI-Rev primer, then PCR-amplified (E2 N-term region) followed by Sanger sequencing (Eurofins Scientific) using *Bsr*GI-For primer (Figure S1, Table S3). To streamline recombinant virus naming and following sequence verification results, rSAV2 and rSAV-patho were renamed rSAV2 E2 VAV, and rSAV2 E2 VAA respectively.

### Western blot analyses of SAV E3E2 furin cleavage

For analysis of SAV E3E2 furin cleavage, 2.5×10^6^ BF-2 cells were seeded in six-well plates. The cells were either mock-infected or infected with SAV2 (SDV, S49P) at an m.o.i. of 1 or 5 and incubated at 15 °C. 72 h after infection the cells were lysed in lysis buffer (150 mM NaCl, 50 mM Tris-HCl pH 8.5, 20 mM EDTA, 1% deoxycholic acid sodium salt, and 1% Triton X-100) with the addition of protease inhibitor cocktail (Roche). For virus purification, wild-type SAV2 was mass produced in BF-2 cells, clarified by low-speed centrifugation at 4,000 rpm for 10 min at 4 °C, then concentrated 10-fold by ultracentrifugation at 25,000 rpm in a SW28 Beckman rotor for 90 minutes at 4 °C and finally purified by ultracentrifugation at 36,000 rpm in a SW41 Beckman rotor for 4 hours at 4 °C on a 25% (w/v) sucrose cushion in TEN buffer (10 mM Tris-HCl pH 7.5, 150 mM NaCl, 1 mM EDTA pH 8). The viral pellet was then resuspended in TEN buffer. Aliquots of cell lysates or sucrose-cushion purified virions were incubated with cracking buffer composed of 20% glycerol, 10% β-mercaptoethanol, 100 mM Tris pH 7, 2% SDS, and bromophenol blue and heated at 100 °C for 5 min. The samples were then separated by SDS-PAGE gel electrophoresis on a 4 to 12% gradient polyacrylamide Bis-Tris gel (NuPage, Invitrogen). After transfer on a polyvinylidene difluoride (PVDF) membrane (Immobilon-P; Millipore), separated proteins were detected using either a mouse monoclonal antibody (mAb 17H23) against SAV E2, or a mAb against SAV E1 (78K5) ^27^, or against the cellular housekeeping protein GAPDH (loading control, Enzo Life Sciences). Immunolabeled proteins were visualized with HRP-conjugated goat anti-mouse antibodies using an enhanced chemiluminescence (ECL) detection system (Pierce), and image acquisitions of Western blot signals were performed using a ChemiDoc Imaging System (Bio-Rad).

### Virus titration and growth kinetics

BF-2 cells were infected with rSAVs at an m.o.i. of 0.1. Supernatants from rSAV-infected cells were collected at 0, 2-, 4-, 7-, 10-, and 14-d.p.i. Ten-fold dilutions of supernatants were used to infect fresh BF-2 cells seeded in 96-well plates. Viral titrations were performed in duplicate experiments. At 7 d.p.i., virus titers were determined by immunofluorescence assays and counting of fluorescent foci as described below (fluorescence focus assay).

### Immunofluorescence and fluorescence focus assays

rSAV-infected BF-2 cells were fixed at 7 d.p.i. with a cold solution of ethanol:acetone (1:1, v/v) for 20 min at -20 °C and then air dried at room temperature. Immunofluorescence assays were performed using a mAb directed against SAV E2 protein (17H23) previously obtained in our laboratory ^27^. Fixed BF-2 cells were incubated for 45 min at room temperature with primary mouse mAb 17H23 diluted 1:10000 in Tween 0.05% (PBS-T). The cells were subsequently washed three times in PBS-T. The cells were then incubated for 45 min at room temperature with Alexa Fluor (AF) 488-conjugated goat anti-mouse antibody (Invitrogen ThermoFisher) diluted 1:800 in PBS-T. The cells were then washed three times in PBS-T. For fluorescence focus assays, positively-infected cell foci were visualized and counted under an Eclipse TE 200 inverted fluorescence microscope (Nikon). Viral titers were calculated taking into account the dilution factors and are expressed as fluorescent focus units per mL (FFU/mL). Immunofluorescence assays were also undertaken at 7-and 10-d.p.i. for microscopy image acquisitions as detailed above with the addition of Hoechst for labeling of cell nuclei in the secondary antibody solution. The immunolabeled cells were imaged using an Axio Observer.Z1 inverted fluorescence microscope with a 10× objective (Zeiss) equipped with a CoolSNAP HQ^2^ CCD camera (Photometrics Roper Scientific). Microscopy image acquisitions were performed using Zen 2.3 software (Zeiss).

## Ethics statement

All *in vivo* experimentations involving rainbow trout strictly adhered to the European guidelines and recommendations on animal experimentation and welfare (European Union Directive 2010/63). All experimental procedures involving rainbow trout were approved by the local ethics committee on animal experimentation (Comité d’éthique appliqué à l’expérimentation animale pour le centre de recherche INRAE de Jouy-en-Josas; COMETHEA INRAE no. 45) and were authorized by the Ministère de l’Éducation nationale, de l’Enseignement supérieur et de la Recherche (APAFIS authorization no. 29801–2021021110262075 v2). The fish facilities in which experiments were carried out were also approved (authorization no. D-78-322-720). To minimize animal suffering and distress, all manipulations were carried out under light anesthesia performed by bath immersion with tricaine (0.005%, Sigma-Aldrich MS-222). During the entire duration of experiments, fish were monitored twice daily for clinical signs and survival. Upon display of typical infection symptoms, animals were humanely euthanized by bath immersion using a lethal dose of tricaine (0.015%, Sigma-Aldrich MS-222).

## *In vivo* analyses of rSAV2 infection in rainbow trout

Two lines of rainbow trout were used for the *in vivo* analyses of infection by rSAV2 variants: the INRAE synthetic line (average weight 2.64 g) and the AP2 isogenic line (average weight 2.95 g) ^36,85^. For each rainbow trout line, groups of 25 virus-free juveniles housed in individual tanks were infected by intraperitoneal injection (i.p.) with 10^4^ FFU/fish of rSAV2 E2 AAV, VAV, VAA and T1-AAA or were mock infected (serum-free GMEM medium). The rSAV2 with AAF, AAD, and AAA triplet motif were not included in the infection trials because of the low virus titers obtained after recovery (Table S4). The fish were kept in separate groups in 30 L tanks at 10 °C. Mortalities were recorded daily over a period of 63 days.

To verify rSAV2 sequence stability during the course of the infection trial, the SAV E2 N-term sequences of the rSAV2 variants isolated from infected AP2 rainbow trout were analyzed. Sampling of head kidney and spleen was performed on fish that had succumbed to infection for each group. The organs (from 14 fishes) were collected, kept individually (not pooled), and homogenized using ceramic beads and the Precellys 24 Touch tissue homogenizer (Bertin Technologies). RNA was extracted using the RNeasy Mini Kit (Qiagen) according to manufacturer’s guidelines. cDNA synthesis and RT-PCR were performed using SuperScript IV reverse transcriptase (Invitrogen) and the SEQ172 and SEQ173 primer pair allowing to amplify a region encompassing the E2 N-term region (Table S3). The PCR-amplified region was sequenced by Sanger sequencing (Eurofins Scientific) using each of the above-mentioned specific primers.

## Statistical analyses

Experimental data were analyzed and plotted using GraphPad Prism (version 7). For the *in vivo* rSAV2 infection trials, cumulative mortalities were plotted as Kaplan-Meier survival curves. Statistical groupings were inferred by performing a Log-rank Mantel-Cox test on the Kaplan-Meier survival data. The following convention was used for describing statistical significance: not significant (n.s.) *p* > 0.05; significant (*) *p* ≤ 0.05; highly significant (**), *p* ≤ 0.01.

