## Supplementary figures and images for "*In silico* reconstruction of a salmonid alphavirus virion reveals distinctive structural and molecular features implicated in virulence *in vivo*"

### Supplemental figure S1

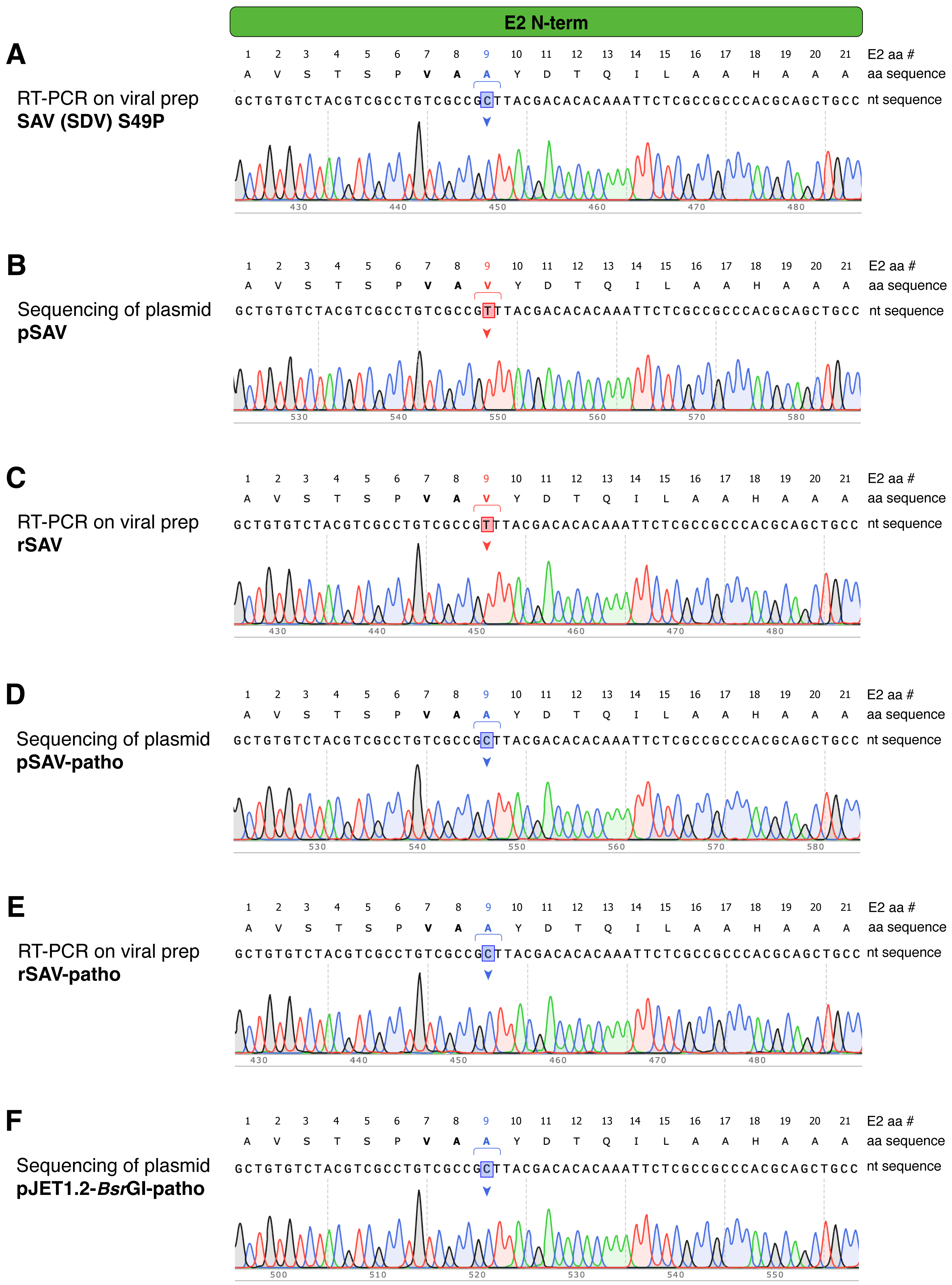

### Supplemental figure S2

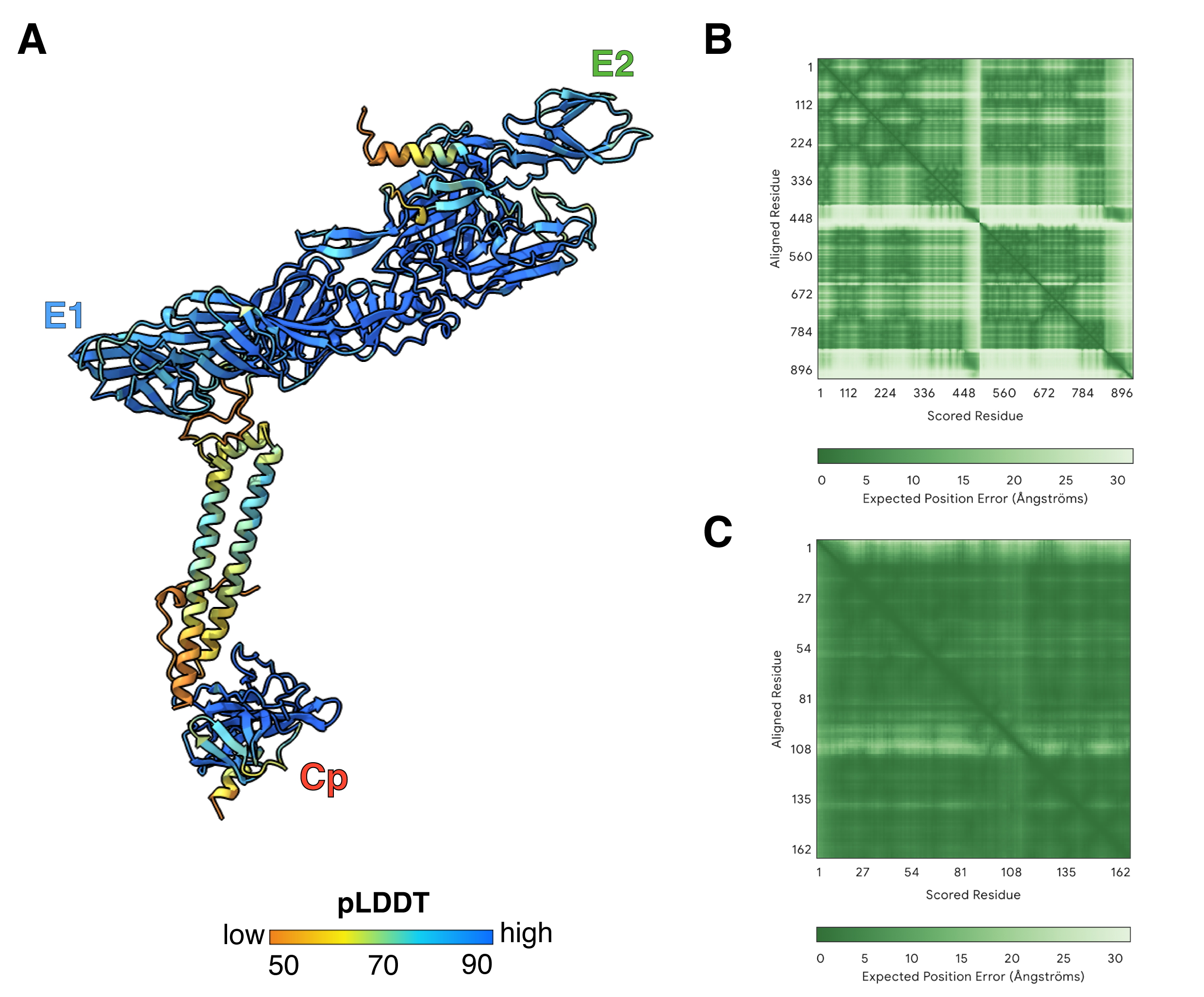

### Supplemental figure S3

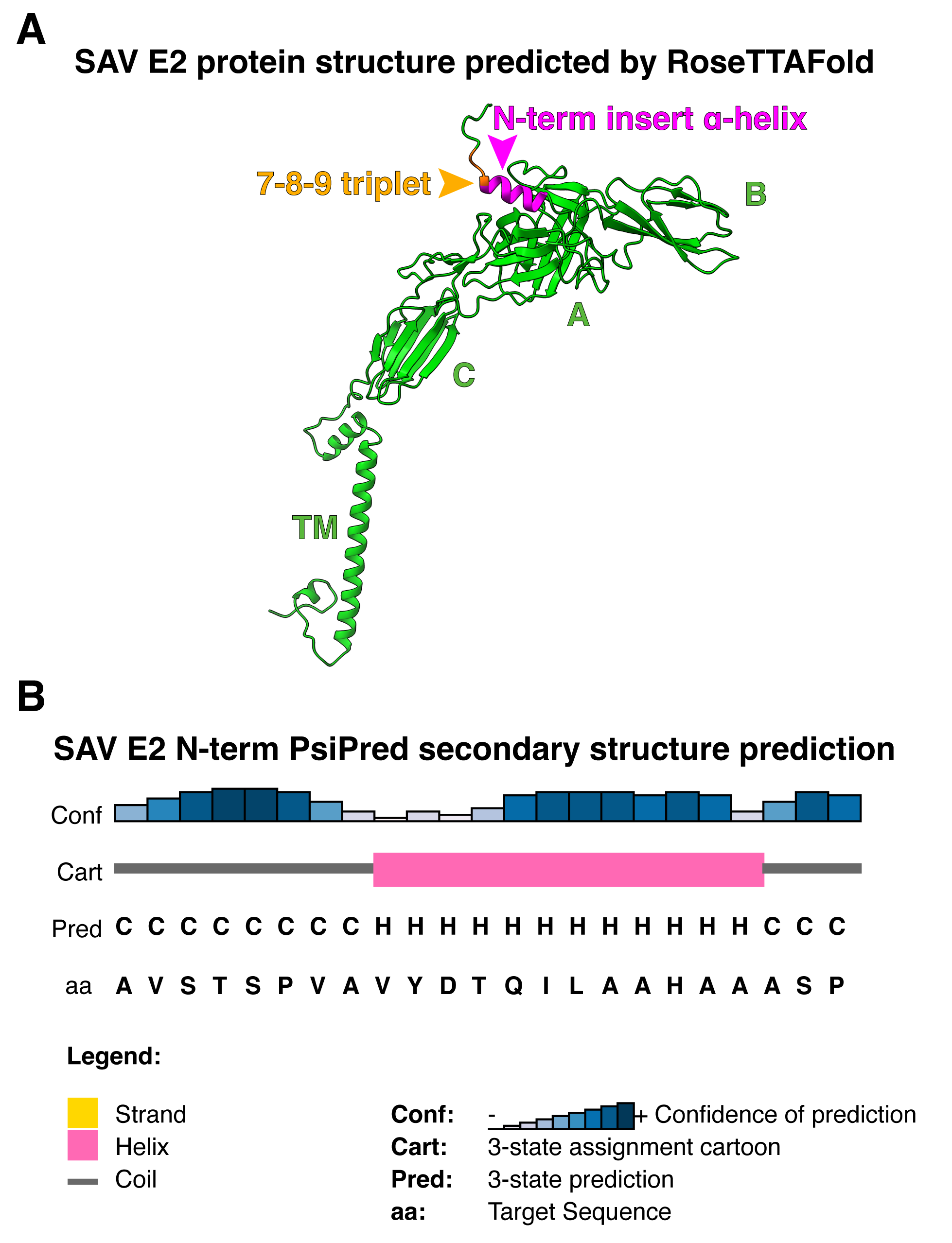

### Supplemental figure S4

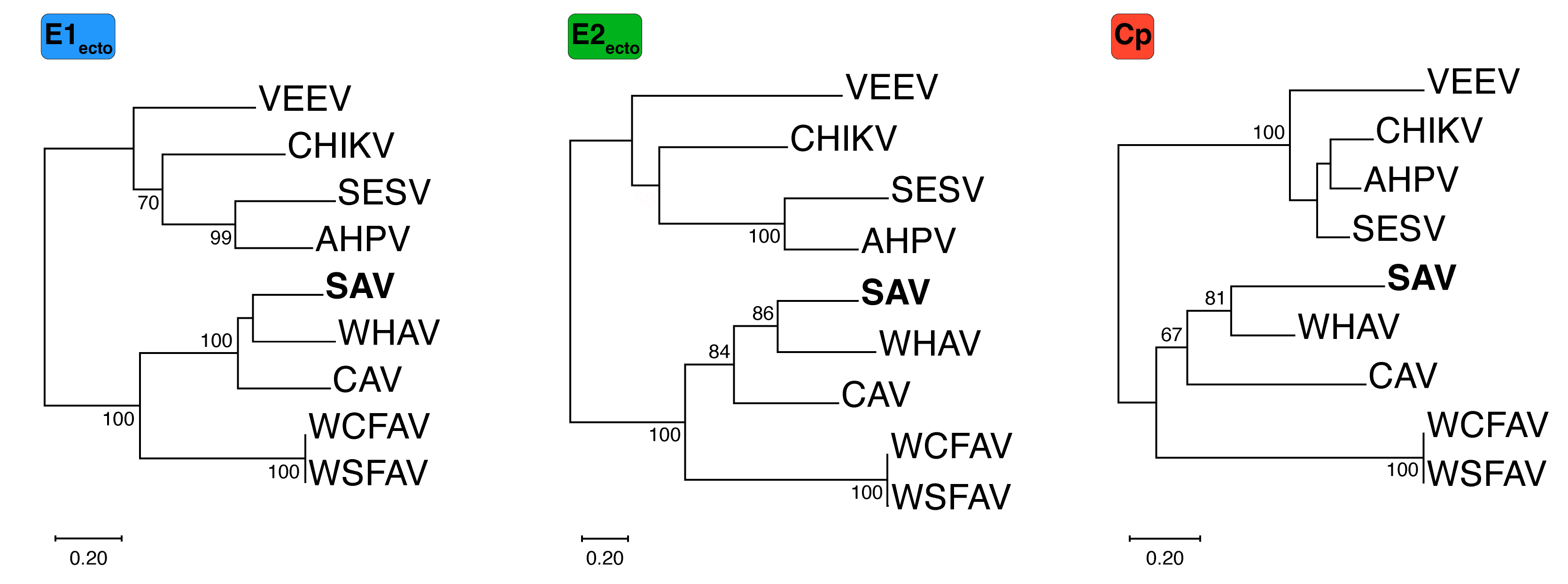

### Supplemental figure S5

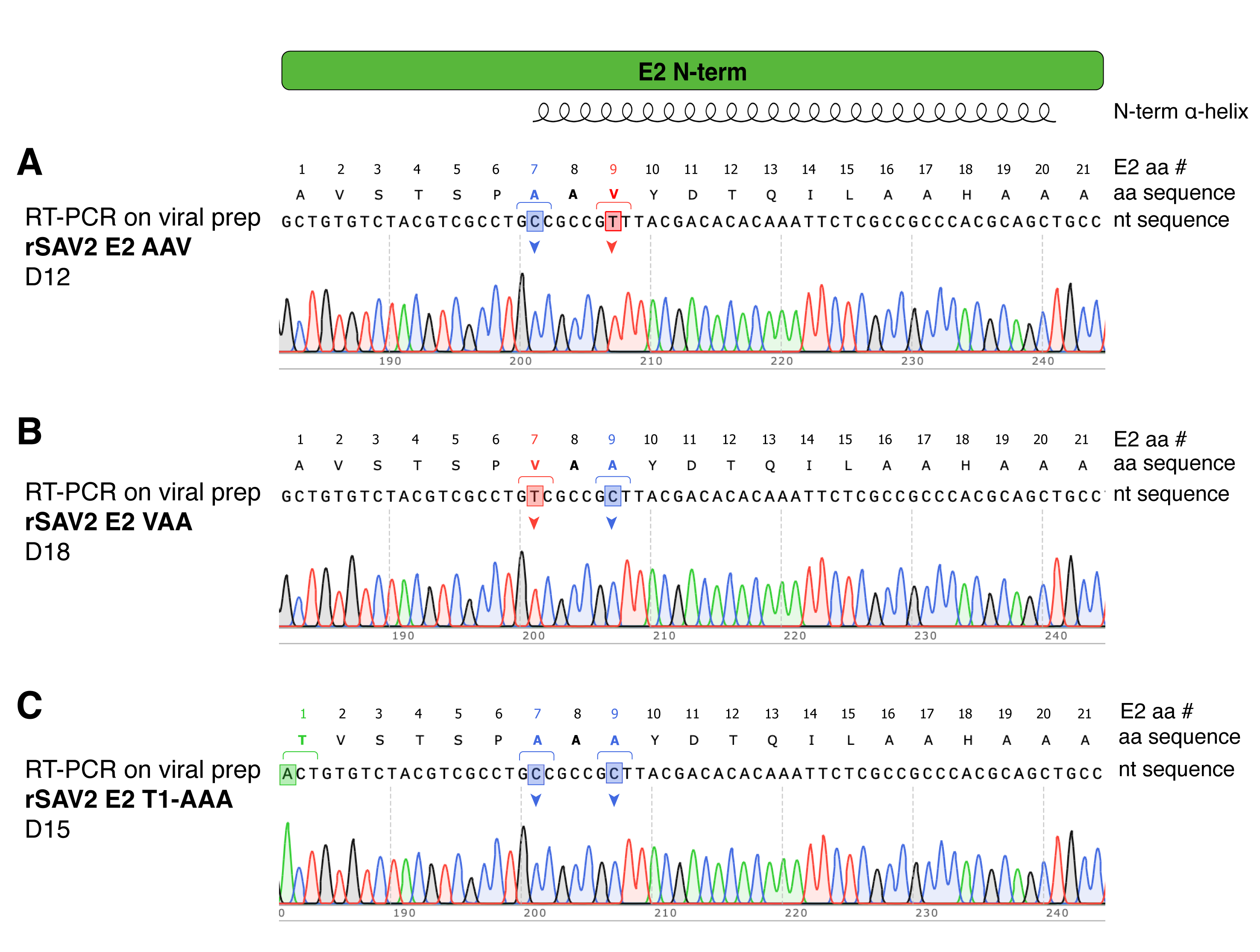

### Supplemental figure S6

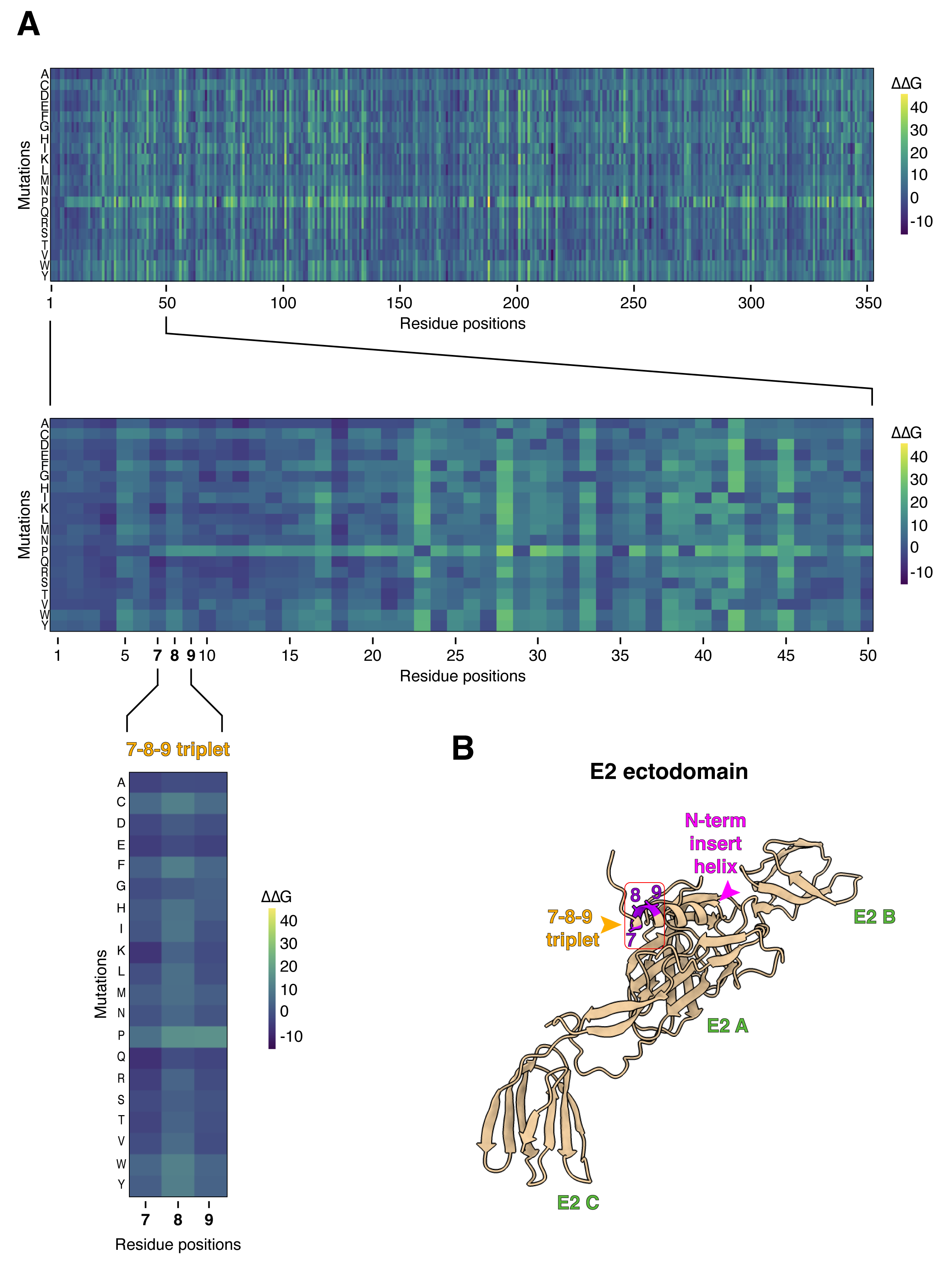
