## Supplemental tables S1-S5 for "*In silico* reconstruction of a salmonid alphavirus virion reveals distinctive structural and molecular features implicated in virulence *in vivo*"

| Alphavirus polyprotein | NCBI accession no. |
|---|---|
| Venezuelan equine encephalitis virus (VEEV) structural polyprotein | AGE98316.1 |
| Eastern equine encephalitis virus (EEEV) structural polyprotein | ABL84687.1 |
| Sindbis virus (SINV) structural polyprotein | AAM10630.1 |
| Western equine encephalitis virus (WEEV) structural polyprotein | AAF28340.1 |
| Eilat virus (EILV) structural polyprotein | AFR68770.1 |
| Ross River virus (RRV) structural polyprotein | ACV66992.1 |
| Semliki Forest virus (SFV) structural polyprotein | NP_463458.1 |
| Mayaro virus (MAYV) structural polyprotein | AAL79764.1 |
| O'nyong-nyong virus (ONNV) structural polyprotein | AAC97205.1 |
| Chikungunya virus (CHIKV) structural polyprotein | NP_690589.2 |
| Southern elephant seal virus (SESV) structural polyprotein | AEJ36233.1 |
| Alaskan harbor porpoise alphavirus (AHPV) structural polyprotein | QJE50388.1 |
| Salmonid alphavirus (SAV) structural polyprotein | CAC87661.1 |
| Wenling hagfish alphavirus (WHAV) structural polyprotein | AVM87430.1 |
| Comber alphavirus (CAV) structural polyprotein | QEP51830.1 |
| Wenling crested flounder alphavirus (WCFAV) structural polyprotein | AVM87428.1 |
| Wenling striated frogfish (WSFAV) structural polyprotein | AVM87426.1 |

**Table S1. Polyprotein sequences used for *Alphavirus* genus phylogenetic analysis.** The alphavirus polyproteins used to generate the genus-wide phylogenetic tree are shown along with the corresponding NBCI accession numbers and hyperlinks.

Supplementary tables

| Subtype | Isolate/strain | Host | NCBI accession no./reference | Raw sequence reads |
|---|---|---|---|---|
| SAV2 | SCO-G524-07 | Rainbow trout | Gallagher et al., 2020 | PRJNA599596 |
| SAV2 | SCO-G582-07 | Rainbow trout | Gallagher et al., 2020 | PRJNA599596 |
| SAV2 | SCO-G399-09 | Rainbow trout | Gallagher et al., 2020 | PRJNA599596 |
| SAV2 | SDV | Rainbow trout | AJ316246.1 | |
| SAV2 | SCO-G573-09 | Common dab | Gallagher et al., 2020 | PRJNA599596 |
| SAV2 | SCO_09_6229_C | Atlantic salmon | MZ395647.1 | |
| SAV2 | SCO-G407-09 | Atlantic salmon | Gallagher et al., 2020 | PRJNA599596 |
| SAV2 | SCO 09 15772 ZBQ | Atlantic salmon | MZ395648.1 | |
| SAV2 | SCO-G572-09 | Common dab | Gallagher et al., 2020 | PRJNA599596 |
| SAV2 | SCO 09 6067 ZAF | Atlantic salmon | MZ395646.1 | |
| SAV2 | SCO-G424-09 | Atlantic salmon | Gallagher et al., 2020 | PRJNA599596 |
| SAV2 | SCO07-4619 | Atlantic salmon | MH708652.1 | |
| SAV2 | SAV2b-BC36 | Atlantic salmon | Macqueen et al., 2021 | |
| SAV2 | SCO-06-17014_F | Atlantic salmon | MZ395649.1 | |
| SAV2 | MR-R5-2011 | Atlantic salmon | MZ395641.1 | |
| SAV2 | SAV2-Nor-1 | Atlantic salmon | KF668081.1 | |
| SAV2 | SAV2-Nor-2 | Atlantic salmon | KF668082.1 | |
| SAV2 | MR-N1-2011 | Atlantic salmon | MZ395642.1 | |
| SAV2 | T-1-2012 | Atlantic salmon | MZ395644.1 | |
| SAV2 | MR-R3-2012 | Atlantic salmon | MZ395645.1 | |
| SAV2 | ST-1-2011 | Atlantic salmon | MZ395643.1 | |
| SAV2 | SAV2a-BC25 | Atlantic salmon | Macqueen et al., 2021 | |
| SAV2 | SAV2a-BC64 | Atlantic salmon | Macqueen et al., 2021 | |
| SAV2 | SAV2b-BC8 | Atlantic salmon | Macqueen et al., 2021 | |
| SAV2 | ALV423 | | Macqueen et al., 2021 | |
| SAV2 | ALV426 | | Macqueen et al., 2021 | |
| SAV3 | SPDV-F08-12 | Atlantic salmon | KF668067.1 | |
| SAV3 | norSAV-SF21-03 | Atlantic salmon | DQ122130.1 | |
| SAV3 | norSAV-Tun1 | | AY604238.1 | |
| SAV3 | SPDV-H07-3 | Atlantic salmon | KF668070.1 | |
| SAV3 | H20-03-2 | Atlantic salmon | MW196361.1 | |
| SAV3 | SPDV-SAVH20-03 | Atlantic salmon | DQ149204.1 | |
| SAV3 | norSAV-R31-04 | Atlantic salmon | DQ122146.1 | |
| SAV3 | SAV3-3-MR-10 | Atlantic salmon | KC122925.1 | |
| SAV3 | SAV3-2-MR-10 | Atlantic salmon | KC122926.1 | |
| SAV3 | SPDV-ST09-7(2) | Atlantic salmon | KF668086.1 | |
| SAV3 | SPDV-N03-8 | Atlantic salmon | KF668075.1 | |
| SAV3 | SPDV-MR07-6(2) | Atlantic salmon | KF668074.1 | |
| SAV3 | SPDV-MR07-6 | Atlantic salmon | KF668073.1 | |
| SAV3 | SPDV-MR07-5 | Atlantic salmon | KF668072.1 | |

| | | | | |
|---|---|---|---|---|
| SAV3 | SPDV-SF07-4(4) | Atlantic salmon | KF668085.1 | |
| SAV3 | SPDV-SF07-4(3) | Atlantic salmon | KF668084.1 | |
| SAV3 | SPDV-SF07-4(2) | Atlantic salmon | KF668083.1 | |
| SAV3 | norSAV-R01-99 | Atlantic salmon | DQ122145.1 | |
| SAV3 | norSAV-SAVF29-03 | Atlantic salmon | DQ122127.1 | |
| SAV3 | SAV3-8-R-10 | Atlantic salmon | KC122921.1 | |
| SAV3 | norSAV-Hav1 | | AY604235.1 | |
| SAV3 | FR18260741 | Atlantic salmon | MN906932.1 | |
| SAV3 | FR12428008 | Atlantic salmon | MN906916.1 | |
| SAV3 | FR12429232 | Atlantic salmon | MN906917.1 | |
| SAV3 | FR16893400 | Atlantic salmon | MN906923.1 | |
| SAV3 | FR14304869 | Atlantic salmon | MN906919.1 | |
| SAV3 | SAV3-7-R-09 | Atlantic salmon | KC122918.1 | |
| SAV3 | SAV3-6-H-10 | Atlantic salmon | KC122922.1 | |
| SAV3 | A140710-13 | Atlantic salmon | JN989318.1 | |
| SAV3 | SAV-Hordaland 2007-3 | Atlantic salmon | KF668079.1 | |
| SAV3 | SAV-Hordaland 2007-2 | Atlantic salmon | KF668078.1 | |
| SAV3 | SAV-Hordaland 2007-1 | Atlantic salmon | KF668077.1 | |
| SAV3 | SPDV Rogaland 3 | Atlantic salmon | KF668076.1 | |
| SAV3 | SPDV-ALV407 | Atlantic salmon | KF668058.1 | |
| SAV3 | SPDV-ALV409 | Atlantic salmon | KF668059.1 | |
| SAV3 | norSAV-PD97.N3 | Rainbow trout | DQ122134.1 | |
| SAV3 | norSAV-PD97.N2 | Atlantic salmon | DQ122133.1 | |
| SAV3 | norSAV-H02-99 | Atlantic salmon | DQ122135.1 | |
| SAV3 | norSAV-N3 | | AY604237.1 | |
| SAV3 | SPDV-H07-1 | Atlantic salmon | KF668069.1 | |
| SAV3 | SPDV-H06-1 | Atlantic salmon | KF668068.1 | |
| SAV1 | SCO-G573-09 | Common dab | Gallagher et al., 2020 | PRJNA599596 |
| SAV1 | SCO-G521-10 | Atlantic salmon | Gallagher et al., 2020 | PRJNA599596 |
| SAV1 | SCO-G524-07 | Rainbow trout | Gallagher et al., 2020 | PRJNA599596 |
| SAV1 | IRE-F10-12 | European plaice | Gallagher et al., 2020 | PRJNA599596 |
| SAV1 | IRE-F7-11 | Common dab | Gallagher et al., 2020 | PRJNA599596 |
| SAV1 | SPDV-ALV418 | Atlantic salmon | KF668061.1 | |
| SAV1 | SPDV-ALV417 | Rainbow trout | KF668060.1 | |
| SAV1 | 4640 | Atlantic salmon | JX163854.1 | |
| SAV1 | SCO-G572-09 | Common dab | Gallagher et al., 2020 | PRJNA599596 |
| SAV1 | F93-125 | Atlantic salmon | AJ316244.1 | |
| SAV1 | SCO-G865-15 | Atlantic salmon | Gallagher et al., 2020 | PRJNA599596 |
| SAV4 | SPDV-ALV290511 | Atlantic salmon | KF668065.1 | |
| SAV4 | SPDV-SCO-29 | | KF668071.1 | |

| | | | | |
|---|---|---|---|---|
| SAV4 | SPDV-ALV416 | Atlantic salmon | KF668057.1 | |
| SAV4 | SAV_04-44 | Atlantic salmon | MH708651.1 | |
| SAV5 | SPDV-ALV230511 | Atlantic salmon | KF668064.1 | |
| SAV5 | SPDV-ALVCHH1 | Atlantic salmon | KF668066.1 | |
| SAV5 | SPDV-ALV160511 | Atlantic salmon | KF668063.1 | |
| SAV5 | SAV_SCO07-192 | Atlantic salmon | MH708653.1 | |
| SAV5 | SAV_SCO10-684 | Common dab | MH341514.1 | |
| SAV5 | SAV_SCO07-4638 | Atlantic salmon | MH708650.1 | |
| SAV5 | IRE-F6-11 | Common dab | Gallagher et al., 2020 | PRJNA599596 |
| SAV6 | F1045-96 | Atlantic salmon | MH238448.1 | |

**Table S2. Nucleotide sequences used in SAV E2 phylogenetic analysis.** For each SAV sequence, the corresponding SAV subtype, isolate or strain, host species, NCBI accession number or source reference and NCBI raw sequence reads number with hyperlinks are provided. Host species: Rainbow trout (*Oncorhynchus mykiss*), Common dab (*Limanda limanda*), Atlantic salmon (*Salmo salar*), European plaice (*Pleuronectes platessa*).

| Primer | Target region & description | F/R | 5' to 3' sequence |
|---|---|---|---|
| SDM64 | SAV E2 N-term, introduces AAF triplet 7-8-9 after mutagenesis | F | TGTGTCTACGTCGCCTGCCGCCTTTT ACGACACACAAATTC |
| SDM65 | SAV E2 N-term, introduces AAD triplet 7-8-9 after mutagenesis | F | TGTGTCTACGTCGCCTGCCGCCGATT ACGACACACAAATTC |
| SDM66 | SAV E2 N-term, introduces AAV triplet 7-8-9 after mutagenesis | F | TGTGTCTACGTCGCCTGCCGCCGTTT ACGACACACAAATTC |
| SDM67 | SAV E2 N-term, introduces AAA triplet 7-8-9 after mutagenesis | F | TGTCTACGTCGCCTGCCGCCGCTTAC GACACACAAATTC |
| SDM68 | SAV E2 N-term, introduces T at position 1 and AAA triplet 7-8-9 after mutagenesis (T1-AAA) | F | GCAGTTCCGCCCGAAAAAGAGGACT GTGTCTACGTCGCCTGCCGCCGCTTA CGACACACAAATTC |
| $\Delta_{4-9}$-For | SAV E2 N-term, for deletion of residues 4 to 9 of E2 (5' phosphorylated) | F | TACGACACACAAATTCTCGCCGCCCA CGCAGCTGCC |
| $\Delta_{4-9}$-Rev | SAV E2 N-term, for deletion of residues 4 to 9 of E2 (5' phosphorylated) | R | AGACACAGCCCTCTTTTTCGGGCGG AACTGCAGGT |
| *Bsr*GI-For | SAV Cp upstream of E2 7-8-9 triplet for sequencing of viral preparations (SAV S49P, rSAV, and rSAV-patho) | F | CATGATGGATGGAGTGTACAATT |
| SEQ13 | SAV Cp upstream of E2 7-8-9 triplet for sequencing (pSAV and pSAV-patho plasmids) | F | GGCGGACATCAAGTTCCAGGTCGCC |
| pJET1.2-For | For sequencing insert sequence cloned in pJET1.2-*Bsr*GI-patho | F | CGACTCACTATAGGGAGAGCGGC |
| *Bsr*GI-Rev | Targets region downstream of E2 sequence, used for RT-PCR | R | CTTGCAGCTCTGTACATACGCAA |
| RT-SEQ172 | SAV E3 upstream of E2 7-8-9 triplet for RT-PCR and sequencing of rSAV2 variants isolated from infected rainbow trout | F | GCGCACCAGCTCTCCTGCTGCTGCCT ATGG |
| RT-SEQ173 | SAV E2 downstream of E2 7-8-9 triplet for RT-PCR | R | CGCGGCGTAAACCTTACCGTCCCTTC CCAGGTATCG |

**Table S3. Primers used in this study.** Shown are the primers used for site-directed mutagenesis, generating a deletion mutant (E2 $\Delta_{4-9}$), sequencing, and RT-PCR. For each primer the target region, description, orientation (Forward, F; Reverse, R), and 5' to 3' sequences are provided.

| Recombinant SAV | Titer (FFU/mL) |
|---|---|
| rSAV2 E2 AAV | $1.5 \times 10^7$ |
| rSAV2 E2 VAV | $1.8 \times 10^8$ |
| rSAV2 E2 VAA | $9 \times 10^7$ |
| rSAV2 E2 AAA | n.d. |
| rSAV2 E2 T1-AAA | $4 \times 10^5$ |
| rSAV2 E2 AAD | $3 \times 10^4$ |
| rSAV2 E2 AAF | n.d. |
| rSAV2 E2 $\Delta_{4\text{-}9}$ | not recovered |

**Table S4. Titers of rSAV2 used in this study.** Shown are the rSAV2 with E2 variations and the corresponding titers that were obtained. n.d. not determined because titer is below the limit of detection of $10^2$ FFU/mL. For rSAV2 E2 Δ4-9, despite several attempts the virus was not recoverable.

| E2 N-term variant | E3E2 junction sequence & furin cleavage site | ProP 1.0 score | PiTou 3 score |
|---|---|---|---|
| AAV | $_{58}$LIAVTTCSSA**RKKR**$_{71}$↓$_{1}$AVSTSP**AAV**YDTQILAAHAA$_{20}$ | 0.883 | 11.6 |
| VAV | $_{58}$LIAVTTCSSA**RKKR**$_{71}$↓$_{1}$AVSTSP**VAV**YDTQILAAHAA$_{20}$ | 0.881 | 11.6 |
| VAA | $_{58}$LIAVTTCSSA**RKKR**$_{71}$↓$_{1}$AVSTSP**VAA**YDTQILAAHAA$_{20}$ | 0.873 | 11.6 |
| AAA | $_{58}$LIAVTTCSSA**RKKR**$_{71}$↓$_{1}$AVSTSP**AAA**YDTQILAAHAA$_{20}$ | 0.876 | 11.6 |
| T1-AAA | $_{58}$LIAVTTCSSA**RKKR**$_{71}$↓$_{1}$**T**VSTSP**AAA**YDTQILAAHAA$_{20}$ | 0.863 | 4.3 |
| AAD | $_{58}$LIAVTTCSSA**RKKR**$_{71}$↓$_{1}$AVSTSP**AAD**YDTQILAAHAA$_{20}$ | 0.868 | 11.6 |
| AAF | $_{58}$LIAVTTCSSA**RKKR**$_{71}$↓$_{1}$AVSTSP**AAF**YDTQILAAHAA$_{20}$ | 0.887 | 11.6 |

**Table S5. Effect SAV E2 N-term variations on predicted furin cleavage at the E3E2 junction.** Predicted furin cleavage scores by ProP 1.0 (https://services.healthtech.dtu.dk/services/ProP-1.0/) and PiTou 3 were obtained by using E3E2 junction sequences with E2 N-term variations (E2 position 1 and triplet 7-8-9 motif). ProP scores obtained for sequences above the 0.5 threshold are considered to be predicted to be cleaved. PiTou scores above 0 are predicted to be cleaved (the higher, the better). The furin cleavage motif at E3 C-term is highlighted in bold and blue, while E2 N-term triplet 7-8-9 motif is in bold. Proteolytic cleavage sites are denoted by the following symbol ↓.
